# Nucleosome repositioning in chronic lymphocytic leukaemia

**DOI:** 10.1101/2022.12.20.518743

**Authors:** Kristan V. Piroeva, Charlotte McDonald, Charalampos Xanthopoulos, Chelsea Fox, Christopher T. Clarkson, Jan-Philipp Mallm, Yevhen Vainshtein, Luminita Ruje, Lara C. Klett, Stephan Stilgenbauer, Daniel Mertens, Efterpi Kostareli, Karsten Rippe, Vladimir B. Teif

## Abstract

The location of nucleosomes in the human genome determines the primary chromatin structure and regulates access to regulatory regions. However, genome-wide information on deregulated nucleosome occupancy and its implications in primary cancer cells is scarce. Here, we performed a systematic comparison of high-resolution nucleosome maps in peripheral-blood B-cells from patients with chronic lymphocytic leukaemia (CLL) and healthy individuals at single base pair resolution. Our investigation uncovered significant changes of both nucleosome positioning and packing in CLL. Globally, the spacing between nucleosomes (the nucleosome repeat length, NRL) was shortened in CLL. This effect was stronger in the more aggressive IGHV-unmutated than IGHV-mutated CLL subtype. Changes in nucleosome occupancy at specific sites were linked to active chromatin remodelling and reduced DNA methylation. Nucleosomes lost or gained in CLL in comparison with non-malignant B-cells marked differential binding of 3D chromatin organisers such as CTCF as well as immune response-related transcription factors, allowing delineating epigenetic mechanisms affected in CLL. Furthermore, patients could be better assigned to CLL subtypes according to nucleosome occupancy at cancer-specific sites than based on DNA methylation or gene expression. Thus, nucleosome positioning constitutes a novel readout to dissect molecular mechanisms of disease progression and to stratify patients. Furthermore, we anticipate that the global nucleosome positioning changes detected in our study, like the reduced NRL, can be exploited for liquid biopsy applications based on cell-free DNA to monitor disease progression.

## Introduction

Chronic lymphocytic leukaemia (CLL) is the most common blood cancer in adults in the Western world. Over the past decade novel therapies against specific targets like Bruton tyrosine kinase have emerged in parallel with an increased understanding of its molecular pathogenesis (Quesada et al. 2013; Ferreira et al. 2014; Nabhan and Rosen 2014; Bosch and Dalla-Favera 2019). Previous genome-wide studies of CLL (epi)genomics and transcriptomics have focused on DNA mutations (Landau et al. 2015; Puente et al. 2015) and the interplay between gene expression, deregulated chromatin features like DNA methylation, histone modifications, chromatin accessibility and transcription factor (TF) binding as well as long range chromatin interactions (Ferreira et al. 2014; Kulis et al. 2015; Queiros et al. 2015; Oakes et al. 2016; Rendeiro et al. 2016; Beekman et al. 2018; Mallm et al. 2019; Vilarrasa-Blasi et al. 2021). However, the primary chromatin structure of CLL with respect to the location of nucleosomes within the genome has not been systematically characterized in patient samples, which is also the case for almost all other tumour entities. Nucleosome positioning affects gene expression by modulating accessibility of TFs to their DNA binding sites as an important part of gene regulation in eukaryotes. Nucleosome maps thus provide insight into the gene regulatory mechanisms in disease and can be used for diagnostics in the new generations of liquid biopsies (Shtumpf et al. 2022). Genome-wide maps of nucleosome positions are usually obtained by digesting the linker DNA with nucleases such as MNase and identifying nucleosome positions from sequencing the DNA associated with the histone octamer core (Teif and Clarkson 2019; Shtumpf et al. 2022). Most previous studies of this kind have investigated genome-wide nucleosome positioning in model systems, human cell lines or non-malignant cells (Schones et al. 2008; Valouev et al. 2011; Gaffney et al. 2012; Kundaje et al. 2012; Diermeier et al. 2014; Ho et al. 2014). Related methods include single cell-based Strand-seq, which is inherently more stochastic and moderately correlates with bulk MNase-seq (Jeong et al. 2022), as well as ATAC-seq that is frequently used for cancer cells and tissues (Grandi et al. 2022). The latter method is well suited to detect changes in positioning of 1-2 nucleosomes near TF binding sites (Beekman et al. 2018; Mallm et al. 2019), but does not provide genome-wide nucleosome maps. On the other hand, the need for precisely mapped nucleosome locations across the whole genome of malignant and non-malignant B-cells is becoming very important in view of the increasing use in clinical diagnostics of cell-free DNA (cfDNA), which contains nucleosome-protected genomic regions (Snyder et al. 2016; Im et al. 2021; Lo et al. 2021; Shtumpf et al. 2022). This need goes beyond blood cancers, because even in the case of solid tumours most of the pieces of cfDNA comes from blood cells and only a small fraction originates from tumour tissues. Here, we exploited our high-throughput sequencing experiments with MNase-assisted histone H3 ChIP-seq (Mallm et al. 2019) at a sequencing depth of more than four billion DNA sequence reads to address the following aims: (i) Derive complete genome-wide nucleosome occupancy maps at base pair resolution and detect individual nucleosome position change in human primary malignant versus non-malignant B-cells (NBCs) from patients with CLL and healthy donors. (ii) Dissect both the repositioning of individual nucleosomes and the collective behaviour of nucleosomes in CLL genome-wide at the global chromatin reorganisation level, e.g., with respect to the average spacing between nucleosomes. (iii) Compare nucleosome occupancy in two clinically important CLL subtypes that have the immunoglobulin heavy chain variable region (IGHV) mutated (M-CLL) or unmutated (U-CLL) and are associated with favourable and poor prognosis, respectively.

## Results

### CLL is characterized by global changes of nucleosome positioning

We determined nucleosome positions in malignant B-cells from CLL patients and NBCs pooled from healthy individuals using high-coverage MNase-assisted histone H3 ChIP-seq (Figure 1A). While individual samples were characterized by a heterogeneous distribution of nucleosome profiles, the averages across groups showed clear differences between non-malignant and malignant cells (Supplemental Figure S1). When 4 billion paired-end reads obtained in our experiments were combined to compare the average nucleosome occupancies in 100-bp bins across the genome in CLL patients versus healthy individuals, the Pearson’s correlation was remarkably high (*r* = 0.95), indicating that only a small fraction of the genome undergoes significant changes of nucleosome occupancy in CLL versus NBCs. Random non-specific regions had very similar profiles of nucleosome occupancy for NBCs and CLL (Supplemental Figure S2A), while regulatory regions showed punctate changes (Figure 1B; Supplemental Figures S2B-D, S3). Such regions typically showed distinct nucleosome occupancy changes in nanodomains of size ∼1-5 kb. In addition, we observed two global genome-wide effects. Firstly, the length of nucleosome-protected DNA fragments was shorter in CLL (Figure 1C). Both CLL subtypes were characterized by shorter DNA fragments than non-malignant controls, with U-CLL particularly enriched with subnucleosomal fragments. Interestingly, cell-free DNA of cancer patients is also known to be enriched with shorter fragments in comparison to healthy individuals (Lo et al. 2021). To study nucleosome repositioning separately from this effect, in the following analysis we applied filtering to consider only DNA fragment sizes between 120-180 bp (Figure 1D, Supplemental Figures S1G, S3). A second global, genome-wide effect was the shortening of the average distance between centres of neighbouring nucleosomes, which is given by the nucleosome repeat length (NRL) (Figure 1E). The NRL decreased from ∼200 bp in NBCs to ∼198 bp in M-CLL to ∼195 bp in U-CLL, which is quite large change, comparable to changes between different cell types during stem cell differentiation (Teif et al. 2012). Furthermore, this effect became stronger inside differentially methylated regions (DMRs), where NRL decreased from 200 bp in NBCs to 196 bp in M-CLL to 193 bp in U-CLL.

**Figure 1.**
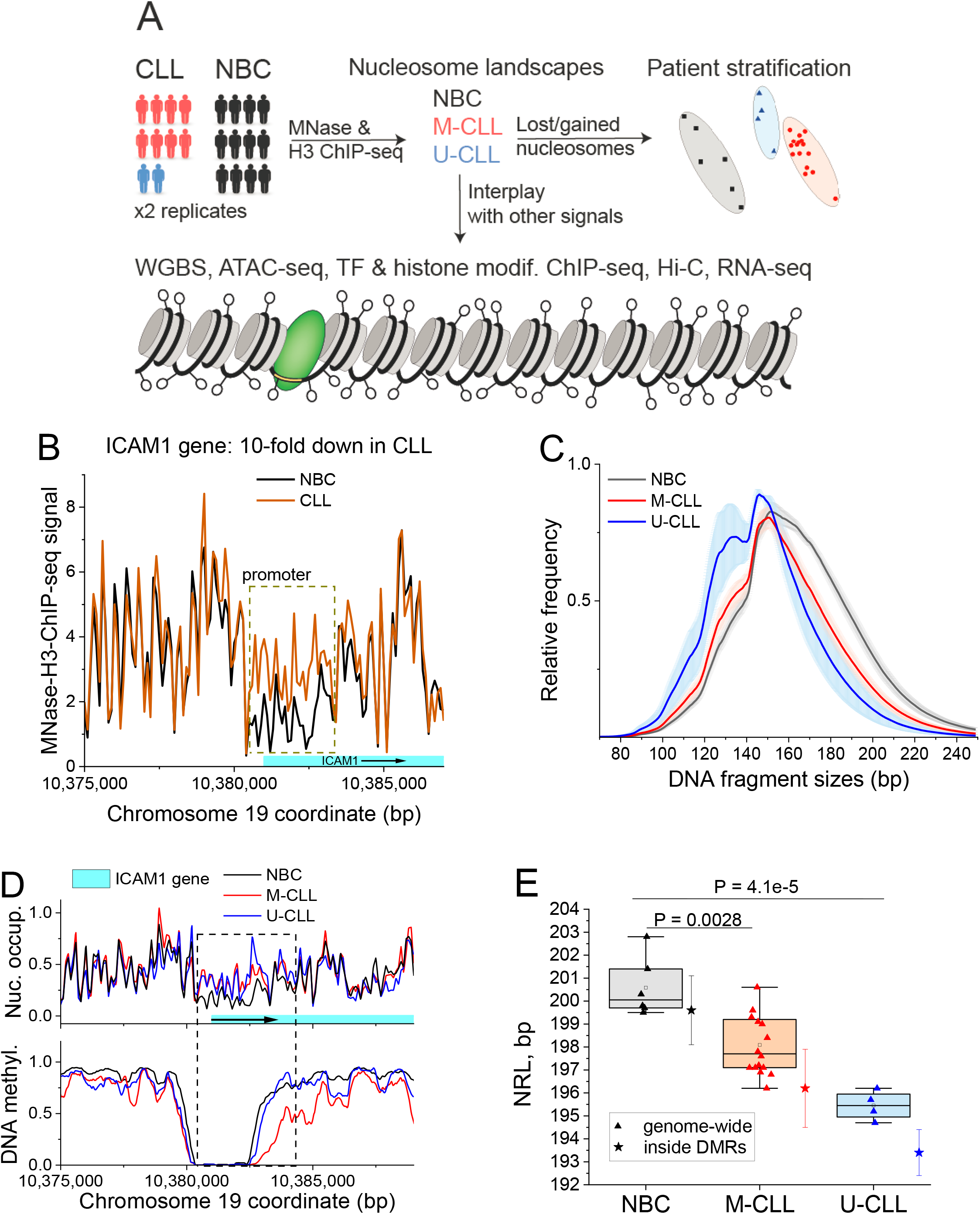
Punctate changes of nucleosome positioning between NBCs, M-CLL and U-CLL. (**A**) Data sets and readouts. Nucleosome landscapes were derived from MNase-assisted H3 ChIP-seq for non-malignant B-cell from healthy donors (NBC) or CLL patients stratified into IGHV mutated (M-CLL) our unmutated (U-CLL). These maps were integrated with data from WGBS, ATAC-seq, ChIP-seq of CTCF, six histone modifications, Hi-C and RNA-seq to dissect molecular mechanisms of nucleosome repositioning. (**B**) Nucleosome occupancy landscapes in NBCs and CLL at a genomic region enclosing promoters of gene ICAM1. Total signal of MNase-assisted H3 ChIP-seq is used without size-selection of DNA fragments. (**C**) Distribution of DNA fragment sizes from MNase-assisted H3 ChIP-seq in NBCs, M-CLL and U-CLL. (**D**) Nucleosome occupancy and DNA methylation for the same region as in panel B but using only 120-180 bp sized DNA fragments. (**E**) Genome-wide nucleosome repeat length (NRL). A decrease from ∼200 bp in NBCs (black) to ∼198 bp in M-CLL (red) to ∼195 bp in U-CLL (blue) is apparent. Each triangle corresponds to one biological sample. Coloured boxes, 25-75% confidence interval; whiskers, range within 1.5 IQR; horizontal line, median, open squares, mean values. Values inside DMRs are shown by star symbols correspond to cohort-averages for NBCs (black), M-CLL (red) and U-CLL (blue) and bars depicting the standard deviation.

### Nucleosome repositioning occurs preferentially in DNA methylation-depleted regions

Next, we asked what distinguishes genomic loci with unchanged or changed nucleosome occupancy in CLL. DNA methylation appeared to stabilise nucleosomes, with nucleosomes in highly methylated regions keeping their locations across NBCs, U-CLL and M-CLL (Figure 2A). In contrast, nucleosomes that shifted more than 20% between NBCs and CLL were mostly depleted of DNA methylation in all conditions (Figure 2B). In addition, we also detected significant differences of nucleosome occupancy associated with CLL-specific changes of DNA methylation (Figure 2C). The difference in nucleosome occupancy was clearly detectable in NBCs versus M-CLL and was even more pronounced in NBCs versus U-CLL. For comparison, we calculated nucleosome profiles around ALU repeats, which are known to stabilize nucleosomes (Teif et al. 2017) (Figure 2D). Nucleosome occupancy around ALUs showed a distinct, stable profile across all conditions, underlining that the variability we detected in CLL cells is coupled to DNA methylation.

**Figure 2.**
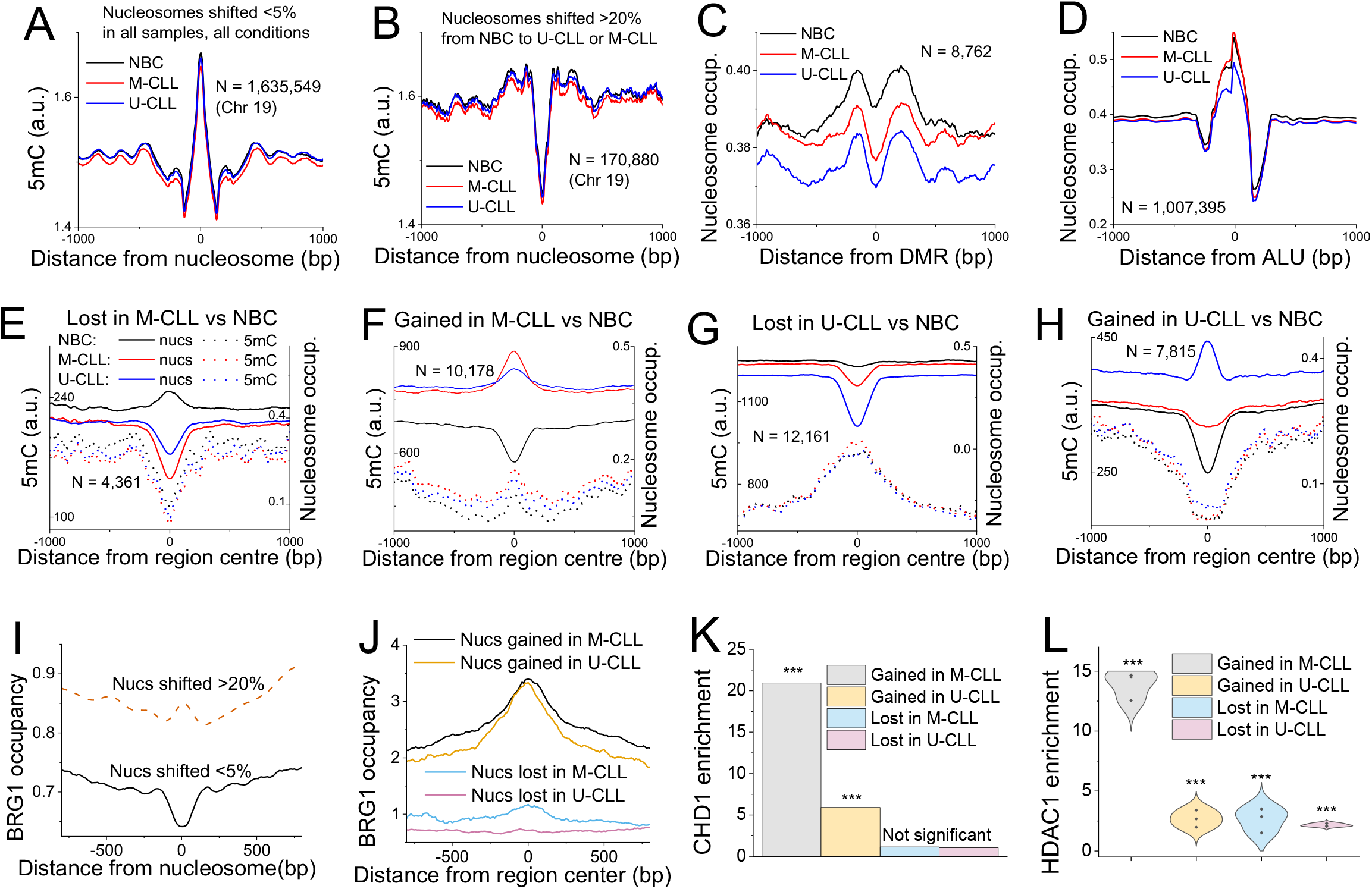
Mechanistic determinants of nucleosome repositioning in CLL. (**A**) DNA methylation occupancy profiles around centres of nucleosome-protected DNA fragments on chromosome 19 for stable nucleosomes with an overlap between different sample types of >95%. (**B**) Same as panel A but for shifted nucleosomes. The overlap of positions between NBCs and CLL was < 80%. (**C**) Nucleosome occupancy around DMRs with decreased DNA methylation in CLL vs NBCs. (**D**) Nucleosome occupancy around ALU repeats. (**E**) Averaged nucleosome occupancy (solid lines) and DNA methylation profiles (dashed lines) around centres of 100-bp regions that lost nucleosomes in M-CLL vs NBCs. The profiles of nucleosome occupancy (solid lines) and DNA methylation (dashed lines) were averaged across all NBCs (black), M-CLL (red) and U-CLL (blue) samples. The number of regions used in the calculation (N) is indicated on the graph. (**F**) Same as panel E but for gained nucleosomes. (**G**) Same as panel E but for lost nucleosomes in U-CLL vs NBCs. (**H**) Same as panel G but for gained nucleosomes. (**I**) BRG1 occupancy in NBCs around centres of stable and shifted nucleosomes at the same regions shown in panel A. (**J**) BRG1 occupancy in NBCs around regions that lost or gained nucleosomes in CLL vs NBCs for the same regions shown in panels E-H. (**K**) Enrichment of CHD1 ChIP-seq peaks determined in lymphoblastoid cell line GM12878 inside regions which undergo nucleosome loss/gain in CLL vs NBCs. (**L**) Enrichment of HDAC1 ChIP-seq peaks determined in peripheral blood mononuclear cells from CLL patients with 11q deletion genotype (GSE216287) inside regions which undergo nucleosome loss/gain in CLL vs NBCs. The violin plots correspond to the distribution based on three ChIP-seq replicates. Fisher test P-value <10e-3 for all points.

### Nucleosome relocation correlates with chromatin remodelling activity

To characterise regions with increased or decreased nucleosome occupancy in CLL in further detail, we segmented the whole genome into non-overlapping 100-bp bins and calculated the normalized nucleosome occupancy for each bin. We then defined regions with stable nucleosome occupancy across all samples from the same group. Stable nucleosome occupancy regions were then compared pairwise between the three conditions. Regions where nucleosome occupancy significantly differed between the two conditions were selected for further analysis and annotated as lost-nucleosome or gained-nucleosome regions correspondingly. This resulted in 4,361 regions that lost nucleosomes in M-CLL vs NBCs, 10,178 regions that gained nucleosomes in M-CLL vs NBCs, 12,161 regions that lost nucleosomes in U-CLL vs NBCs and 7,815 regions that gained nucleosomes in U-CLL vs NBCs (Figure 2E-F). The pairwise comparison of M-CLL and U-CLL revealed 5,117 regions with higher and 307 regions with lower nucleosome occupancy in U-CLL. Figures 2E-G show average profiles of nucleosome occupancy around regions that lost or gained nucleosomes in the two CLL subtypes versus NBCs. The nucleosome occupancy profiles of these regions were more similar between the two CLL subtypes and distinct from NBCs. DNA methylation profiles of these regions showed different behaviour between the two CLL subtypes: regions that lost nucleosomes in U-CLL vs NBCs were enriched with DNA methylation, while those that lost nucleosomes in M-CLL vs NBCs were depleted of DNA methylation. Similarly, regions that gained nucleosomes in U-CLL vs NBCs (but not M-CLL vs NBCs) were depleted of DNA methylation. These DNA methylation profiles were highly reproducible across all samples with the same condition (Supplemental Figure S4). To check whether nucleosome repositioning in CLL is an active process driven by remodellers, we compared the nucleosome maps with the binding of a remodeller BRG1 in NBCs assessed previously by ChIP-seq (Abraham et al. 2013). Nucleosomes that did not change their locations between conditions were depleted of BRG1 in NBCs, whereas shifted nucleosomes (>20% change of their start/end coordinates) were enriched with BRG1 (Figure 2I). The regions that gained nucleosomes in CLL versus NBCs had more than a 3-fold enrichment of BRG1 in comparison with neighbouring regions (Figure 2J). The loci bound by another chromatin remodeller CHD1 in lymphoblastoid B-cells were also enriched in these regions, in particular for gained-nucleosomes in M-CLL with >20-fold enrichment (Figure 2K). Furthermore, the regions that gained nucleosomes in M-CLL also had the largest overlap with loci identified in a recently published ChP-seq dataset as bound by histone deacetylase HDAC1 in peripheral blood mononuclear cells from CLL patients (up to 15-fold enrichment, Figure 2L). This suggests that nucleosome relocation in CLL is subtype-specific and the result of active chromatin remodelling.

### Nucleosome repositioning marks CLL-specific gene regulation

Regions with gained nucleosome occupancy in one of the CLL subtypes vs NBCs were enriched with promoters, active enhancers, CpG islands and CTCF sites, while the most pronounced nucleosome loss was at active CLL enhancers (Figure 3A and B). For CTCF sites, the fold-enrichment increased with the increase of the similarity to the consensus CTCF motif. Regions with gained nucleosome occupancy in CLL were enriched with cancer-related pathways including both generic pathways typically deregulated in cancer as well as CLL-specific pathways such as B-cell receptor signalling (BCR) (Figure 3C and D). Interestingly, the largest group of genes marked by regions that gained nucleosomes in CLL was related to immune system (>350 genes, P=0.01). A similar pairwise comparison of regions with differential nucleosome occupancy between U-CLL and M-CLL is shown in Supplemental Figure S5C-D. In addition, we repeated this analysis including all DNA fragments from MNase-assisted H3 ChIP-seq without size-selection (Supplemental Figure S6). This revealed that the largest fraction of nucleosomes repositioned in CLL versus NBCs resides in 700,000 regions of 100-bp size. These loci were termed “ variable” because their nucleosome occupancy significantly varied across CLL patients while being stable across all NBC samples (Supplemental Figure S6A). The most informative group of nucleosome changes that distinguished CLL was the fraction of gained nucleosomes residing in regions where nucleosome occupancy in CLL increased in comparison with NBCs (Supplemental Figure S6D). The intersection of such gained-nucleosome regions with gene promoters marked the B-cell receptor signalling pathway (BCR) as the top hit (*p* = 1.6e-5), followed by the T-cell receptor signalling pathway (*p* = 3.2e-4) (Supplementary Tables S1-3).

**Figure 3.**
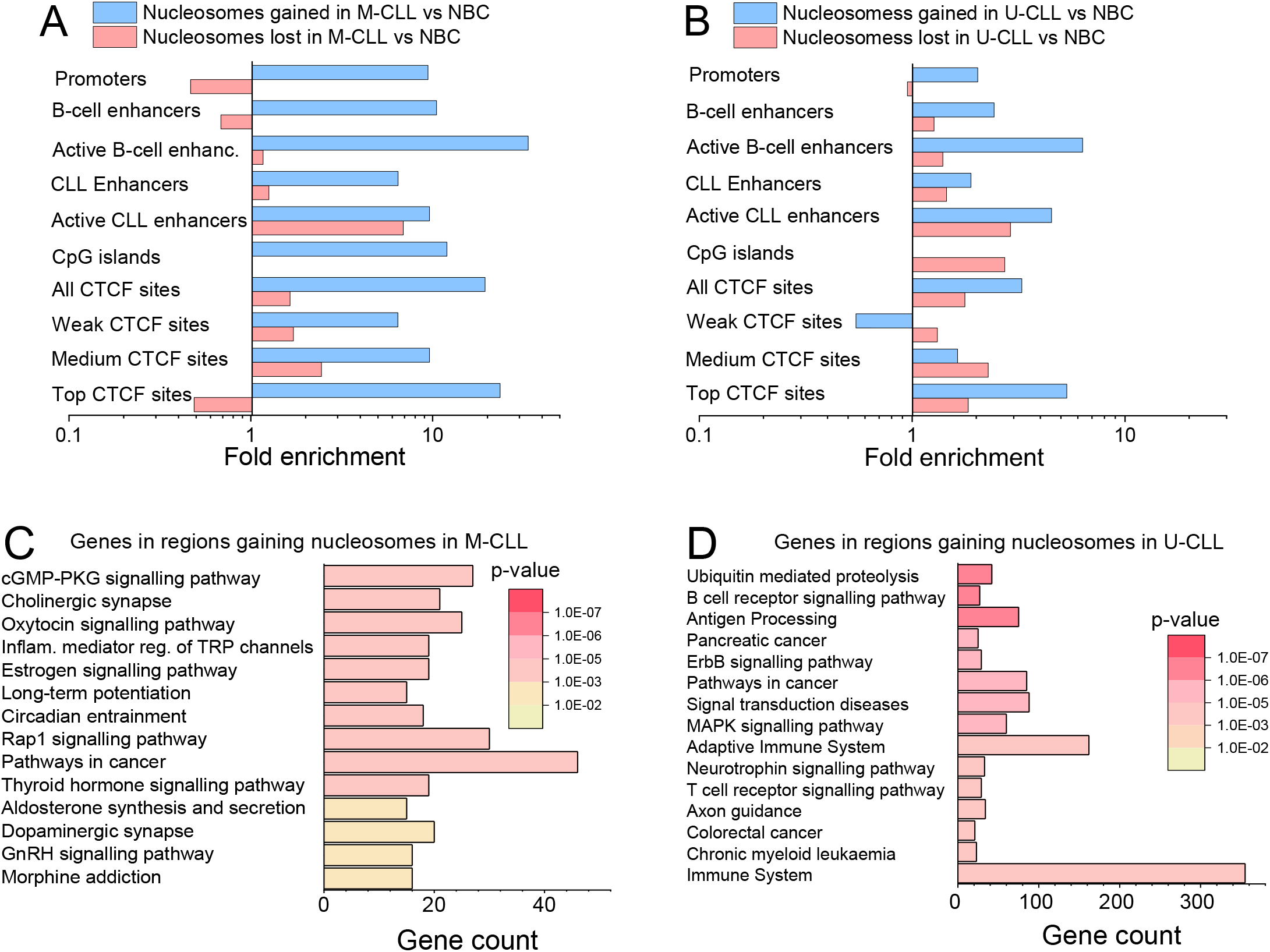
Annotation of sites with differential nucleosome occupancy. (**A**) Genomic regions which lost (red) or gained (blue) nucleosomes in M-CLL vs. NBC. (**B**) Same as panel A but for U-CLL vs. NBC. (**C**) Gene ontology analysis of genes that overlap with regions that gained nucleosomes in M-CLL vs. NBC. (**D**) Same as panel C but for U-CLL vs. NBC.

The correlated differences of nucleosome occupancy and gene activity could also be related to a differential activity of cis-regulatory enhancer regions in NBCs and CLL. Active enhancers (defined previously from multi-omics profiling in this patient cohort (Mallm et al. 2019)) were characterized by nucleosome depletion, whereas all enhancers, which are mostly inactive in each given state, were characterized on average by increased nucleosome occupancy (Figure 4A-D). Of note, active NBC enhancers had very similar DNA methylation profiles between U-CLL and M-CLL, whereas the nucleosome occupancy profiles allowed distinguishing the CLL subtypes (Figure 4C). Nucleosome occupancy differed between CLL and NBCs even at the scale of ∼100,000 bp based on A and B chromatin compartments defined with Hi-C (Vilarrasa-Blasi et al. 2021). Chromatin domains that are open and active in NBCs (A compartments) were characterized by nucleosome depletion, which disappears in U-CLL (Figure 4E). On the other hand, the inactive and closed B compartments displayed a flat profile of nucleosome occupancy in all cell states (Figure 4F). Thus, functionally relevant differences in CLL nucleosome occupancy occurred both at the level of promoters and enhancers, as well as on the mesoscale of active chromatin subcompartments.

**Figure 4.**
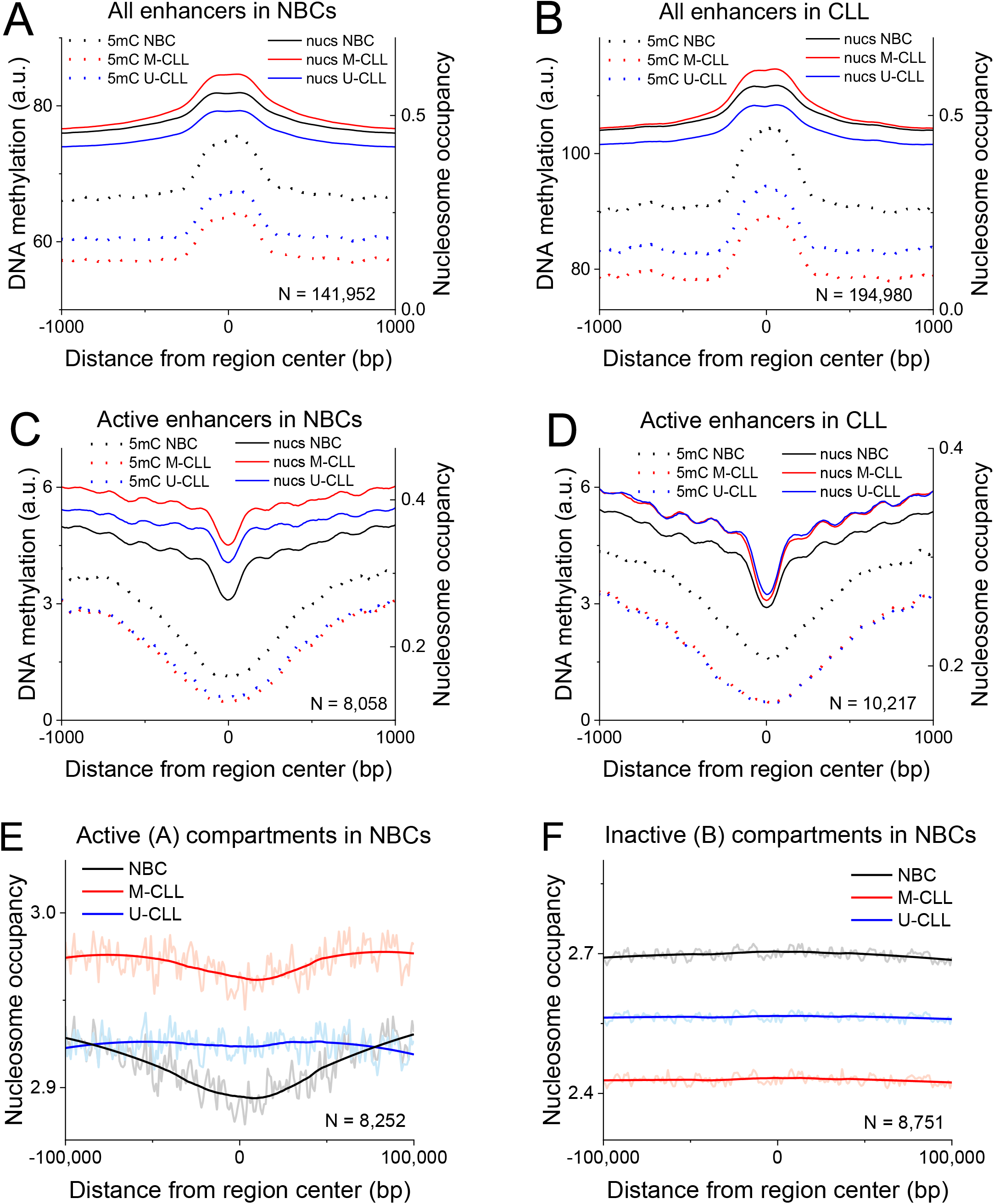
Average nucleosome occupancy and DNA methylation occupancy profiles at cis regulatory elements. (**A**) Averaged nucleosome and DNA methylation occupancy profiles at enhancers in NBCs. (**B**) Same as panel A but for CLL. (**C**) Same as panel A but for active NBC enhancers as determined from ATAC-seq. (**D**) Same as panel C but for active CLL enhancer. (**E**) Averaged nucleosome occupancy profiles within the active A chromatin compartments determined from Hi-C in NBCs by Vilarrasa-Blasi et al., 2021. The profiles are averaged over all NBCs (black), M-CLL (red) and U-CLL (blue) samples. The number of regions (N) is indicated on the graphs. (**F**) Same as panel F but for the in active B compartment.

### Nucleosome positioning, CTCF binding and histone modifications are correlated

Next, we focused on regions with deregulated CTCF binding (Figure 5). Sites that were bound by CTCF both in NBCs and CLL were defined as “ common”. Nucleosome occupancy profiles around common CTCF sites had the same shape for NBCs and CLL and showed a pronounced depletion in the center surrounded by characteristic oscillations reflecting the regular nucleosome array positioned by CTCF. The average DNA methylation profiles were depleted in a wider area +/-200 bp surrounding common CTCF sites both in NBCs and CLL (Figure 5A). In the case of sites that lost CTCF binding in CLL (termed “ lost”), NBCs were characterized by nucleosome depletion around CTCF, but both U-CLL and M-CLL had a peak of nucleosome occupancy instead (Figure 5B). The DNA methylation profiles also gained peaks near lost CTCF sites in CLL. Interestingly, M-CLL and U-CLL displayed different trends: M-CLL had higher nucleosome occupancy around both common and lost CTCF sites. On the other hand, the increase in DNA methylation near lost CTCF sites was higher in U-CLL. We also investigated the effect of CTCF loss in CLL on different histone modifications (Figure 5C). The largest changes were observed for the active enhancer mark H3K4me1, which went down in CLL, as well as the repressive heterochromatin marks H3K9me3 and H3K27me3, which both went up in CLL. In addition, we evaluated a recently reported pan-cancer set of CTCF sites undergoing consistent changes across several cancer types (Fang et al. 2020) (Supplemental Figure S7A-B). Nucleosome occupancy at these CTCF sites was not significantly different between NBCs and CLL, which means that the effect observed in Figure 5 is specific for CLL.

**Figure 5.**
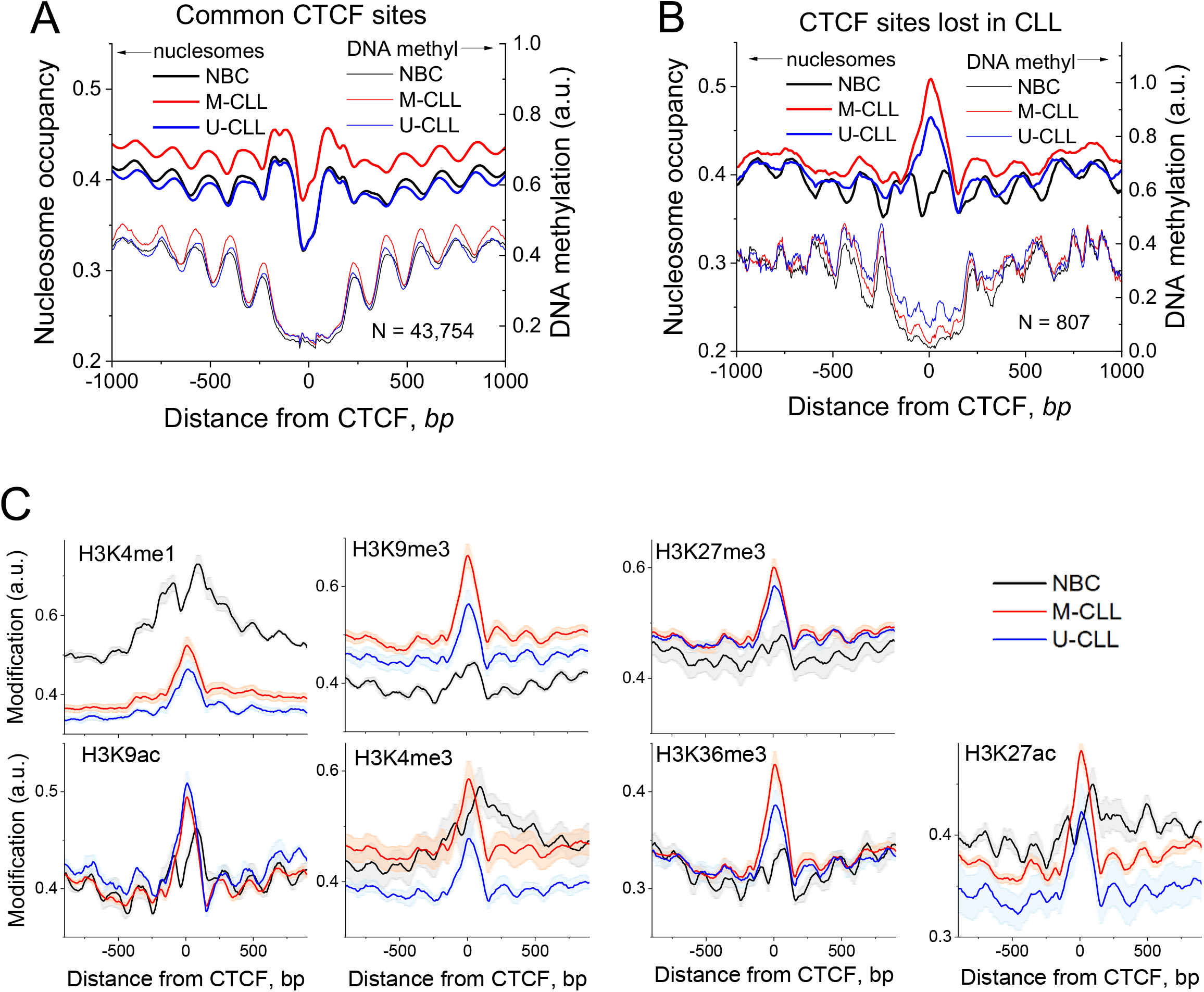
Nucleosome occupancy, histone modifications and DNA methylation around CTCF sites. (**A**) Averaged DNA methylation and nucleosomes occupancy profiles around CTCF motifs which are bound by CTCF both in NBC and CLL. (**B**) Same as panel A but for sits bound by CTCF in NBC but not in CLL. (**C**) Histone modifications around the same set of lost CTCF sites depicted in panel B. The profiles were averaged over all NBCs (black), M-CLL (red) and U-CLL (blue) samples. Standard errors are shown for each line in light colours. The number of regions (N) is indicated on the graph

### Nucleosome repositioning reveals differential TF binding in CLL

To study effects of TF-nucleosome interaction we determined enriched TF motifs inside regions with differential nucleosome occupancy in CLL (Figure 6A), and calculated nucleosome profiles around their binding sites (Figure 6B-C). This analysis revealed three major cases: (i) For one set of TF binding sites, nucleosome profiles in NBCs, M-CLL and U-CLL had a similar shape but differed in average nucleosome occupancy (e.g., ZNF263, PAX9 and PAX5 in Figure 6B). For these motifs, U-CLL and M-CLL were more similar to each other than to NBCs. (ii) In NBCs, M-CLL and U-CLL the shape of nucleosome profiles was similar but NBCs and M-CLL profiles located closer to each other and further away from U-CLL (e.g., TCLF5, HES5 and E2F3 in Figure 6B). In this group, the different average nucleosome occupancy level likely reflects the change in local nucleosome landscape not related directly to the binding of a given TF but rather determined by the binding of other factors nearby. (iii) The shape of nucleosome occupancy profiles changed between CLL and NBCs. These included the transition from low nucleosome occupancy in NBCs to a peak in CLL, e.g., JUND, PKNOX2, FOSL1 (Figure 6C). Such nucleosome profile changes were accompanied by gene expression differences in CLL versus NBCs, e.g., a ∼2-fold reduction for FOSL1. Supplemental Tables S4 and S5 show nucleosome occupancy profiles of TFs enriched in regions with differential nucleosome occupancy defined above but limited to binding sites inside CLL-specific ATAC-seq peaks.

**Figure 6.**
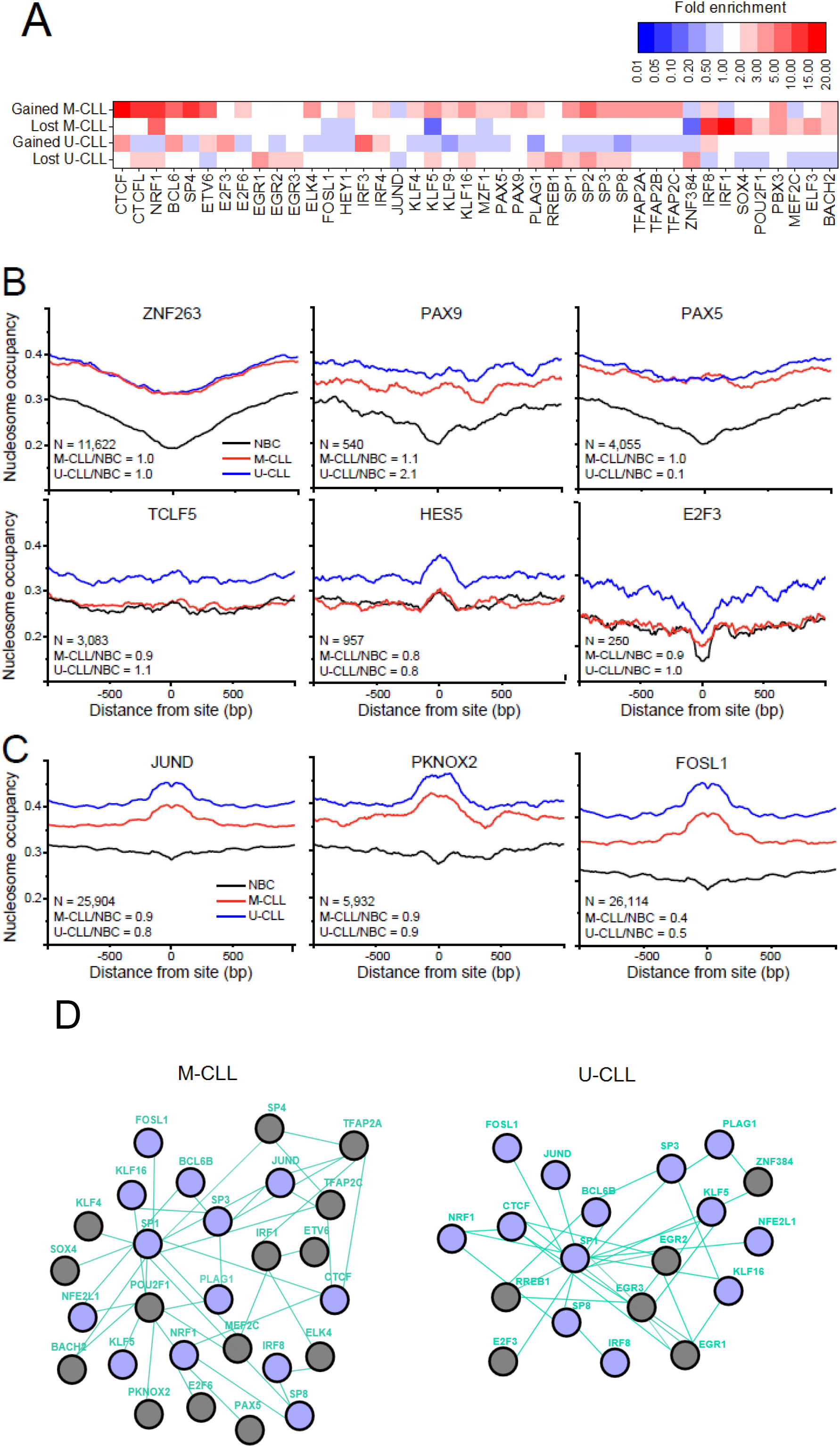
Analysis of TF binding in regions with differential nucleosome occupancy. (**A**) Heatmap showing fold-enrichment levels TFs with >2-fold enrichment of their binding sites in regions that lost or gained nucleosomes in M-CLL or U-CLL versus NBCs, or significantly changed shape of their binding profile in CLL. (**B**) Nucleosome occupancy profiles around binding sites of TFs inside regions that lost ATAC-seq signal in CLL vs NBCs. (**C**) Nucleosome occupancy profile around binding sites of TFs inside regions that gained ATAC-seq signal in CLL vs NBCs. The profiles are averaged over all NBC (black), M-CLL (red) and U-CLL (blue) samples. The number of regions (N) and fold expression change of the corresponding TF in two conditions – M-CLL versus NBC and U-CLL versus NBC, are indicated on the graph. (**D**) Nucleosome repositioning marks distinct TF pathways, which are different between M-CLL and U-CLL while some network features remain common. Green TF connecting lines, known relationships of type “ controls expression” based on pathwaycommons.org; purple circles, TFs affected both in U-CLL and M-CLL; grey circles, TFs unique for CLL subtypes.

In addition to the analysis based on nucleosome-size DNA fragments above, we also performed a similar analysis for all DNA fragments without size-filtering. The analysis of 2,508 promoters with gained-nucleosomes revealed B-cell receptor signalling as the top enriched pathway (Supplemental Tables S1-2) and the so-called CG-box as the top enriched motif (p-value = 1.1e-37, 591 out of 2,508 sites, Supplemental Table S3), suggesting that TFs recognizing this motif such as SP1/2 and EGR1/2 can be involved in CLL deregulation. The expression of genes encoding these TFs was only moderately changed in comparison with non-malignant controls (log2 fold change -0.65 for SP1, -2.4 for SP2, -3.0 for EGR1, - 0.26 for EGR2), indicating that nucleosome repositioning can be more informative than gene expression in peripheral blood B-cells for assessing TF activity. Supplemental Figure S7C-D shows that nucleosome occupancy profiles around SP1 change from nucleosome depletion across all SP1 sites to nucleosome occupancy peak for a subset of SP1 sites covered by “ variable” nucleosomes in CLL. A less frequent motif present in promoters of these genes can be recognized by SOX11 and SOX4 among other TFs (Supplemental Table S3). SOX11 was proposed as a prognostic marker in CLL (Roisman et al. 2015), while SOX4 plays a role in other B-cell malignancies (Sarkar and Hochedlinger 2013). Interestingly, SOX4 was 128-fold down-regulated in CLL vs. non-malignant controls based on our RNA-seq data. A SP1 binding site inside SOX4 promoter was associated with a distinct nucleosome peak appearing at the place of the nucleosome depletion in non-malignant controls (Supplemental Figure S2D), suggesting that differential SP1/2 or EGR1/2 binding might occur upstream of SOX4 deregulation.

### CLL patients can be stratified based on nucleosome occupancy

We examined the classification of samples based on different epigenetic parameters. First, we asked whether it is possible to distinguish NBCs, M-CLL and U-CLL based on gene expression. Figure 7A shows that principal component analysis (PCA) of gene expression values clearly distinguished CLL from NBCs, but not the CLL subtypes from each other. Second, we attempted the same classification using DNA methylation of gene promoters. Figure 7B depicts the clear differences in DNA methylation between sample types. However, promoter DNA-methylation did not segregate the three groups very effectively and there was a partial overlap among the three clouds/sample types. Finally, we asked whether the nucleosome repositioning as identified in this study can serve as a novel biomarker. For example, nucleosome occupancy at the 100-bp regions which lost nucleosomes in U-CLL vs NBCs in each of the 26 samples (including 6 NBCs, 4 U-CLL and 16 M-CLL) was informative in PCA. As shown in Figure 7C-E, PCA based on nucleosome occupancy allows distinguishing not only CLL from NBCs, but also M-CLL from U-CLL. Notably, regions with differential nucleosome occupancy defined using only one CLL subtype worked to distinguish the other CLL subtype from NBCs as well (Figure 7C-F). Interestingly, regions that gained or lost nucleosomes in U-CLL vs NBCs or lost nucleosomes in M-CLL vs NBCs allowed good stratification of all studied subtypes (Figure 7C-E), whereas regions that gained nucleosomes in M-CLL were common between CLL subtypes and less effective in stratifying patients (Figure 7F). Our additional calculations showed that unlike gained-nucleosome and lost-nucleosome regions, nucleosome occupancy or DNA methylation inside other genomic features such as promoters, enhancers or CTCF sites is not as effective in distinguishing M-CLL, U-CLL and NBCs (Supplemental Figures S8-S9).

**Figure 7.**
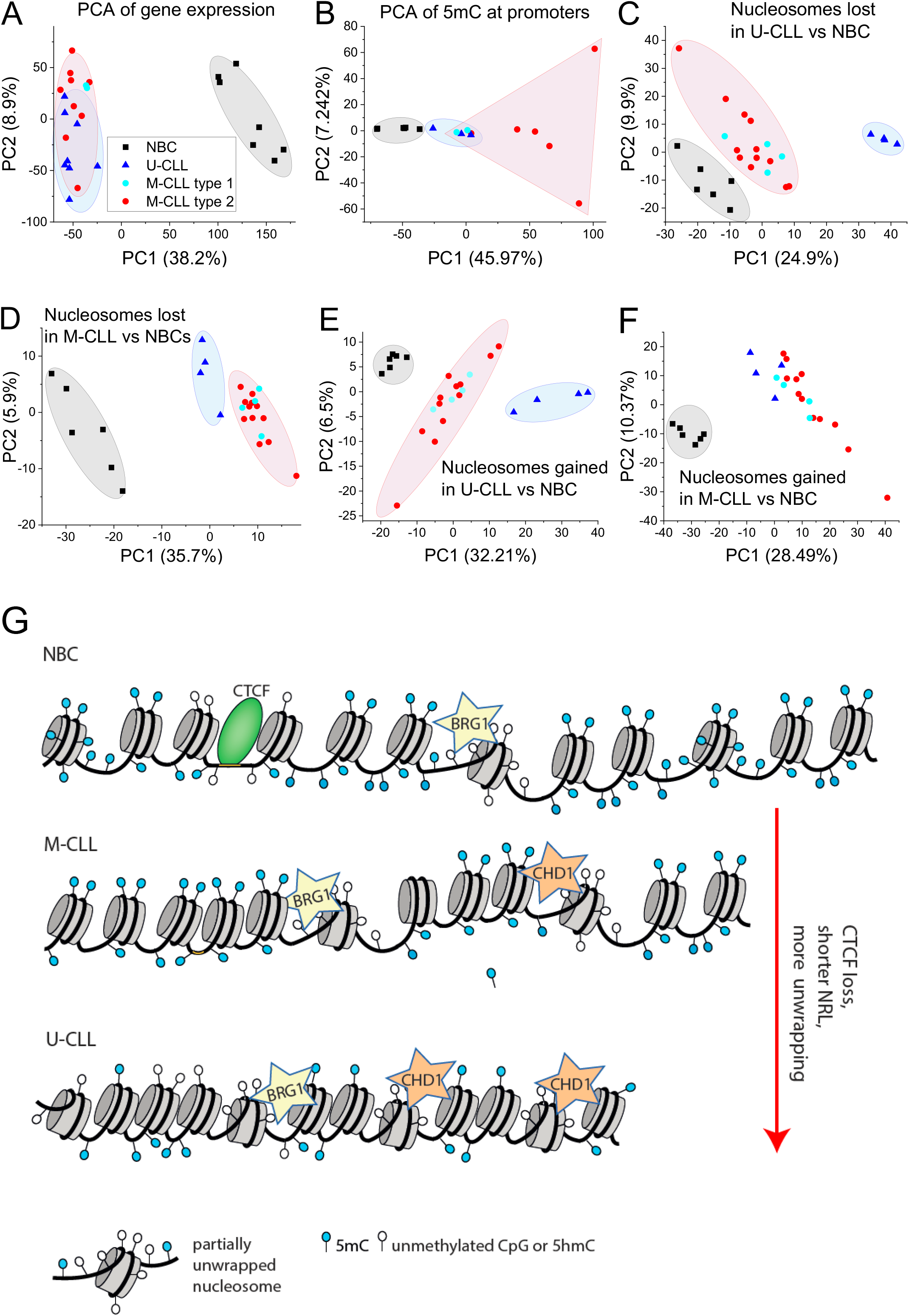
Nucleosome positioning as a new marker to stratify CLL patients. (**A-F**) Principal component analysis (PCA) based on gene expression, DNA methylation and nucleosome occupancy. Each dot represents a replicate from NBC (black squares), M-CLL (red circles) and U-CLL (blue triangles) (**A**) PCA based on gene expression. (**B**) PCA based on 5mC at promoters. (**C**) PCA based on nucleosome occupancy at regions with reduced nucleosome occupancy in U-CLL vs NBC. (**D**) Same as in panel C but for regions with reduced nucleosome occupancy in M-CLL vs NBC. (**E**) Same as panel C but for regions with increased nucleosome occupancy. (**F**) Same as panel D but for regions with increased nucleosome occupancy. (**G**) A scheme of molecular mechanisms of nucleosome repositioning in CLL. CLL B-cells have shorter NRL and more partially unwrapped nucleosomes than NBCs. These features are linked to differential DNA methylation and action of chromatin remodellers. The aberrant nucleosome positioning in CLL is more pronounced for the more aggressive U-CLL subtype as compared to M-CLL.

## Discussion

In CLL, large progress in understanding the contribution of chromatin features to disease pathogenesis has been made in recent years (Beekman et al. 2018; Gaiti et al. 2019; Mallm et al. 2019; Pastore et al. 2019; Rendeiro et al. 2020; Vilarrasa-Blasi et al. 2021). However, the contribution of nucleosome positioning has not been integrated. This essential knowledge gap was addressed here by compiling occupancy maps at single nucleosome resolution in NBCs, U-CLL and M-CLL. Based on these data, our study introduces nucleosome positioning as a powerful biomarker that reveals novel mechanistic insights of chromatin-mediated deregulation in CLL (Figure 7G). We detected changes of the nucleosome landscape at two levels. Firstly, our analysis revealed global genome-wide changes with a striking decrease of the NRL (Figure 1E). Furthermore, the distribution of nucleosome-protected DNA fragments was skewed towards subnucleosomal fragments in the case of M-CLL, and even more so for U-CLL, while NBCs were enriched with nucleosome-plus-linker particles. This finding has important implications for understanding nucleosome positioning patterns in malignant and non-malignant B-cells for cancer diagnostics based on biopsy approaches that analyse the size distribution of cell-free DNA (cfDNA) from body fluids as a tumour marker (Lo et al. 2021; Shtumpf et al. 2022). NRL changes at the scale of few bp are known to occur during cell differentiation (Teif et al. 2012), but have not been evaluated previously in paired normal/cancer cells of the same type. Importantly, NRL changes reported here were visible not only between CLL and NBCs, but also between the two subtypes of CLL, with the more aggressive U-CLL subgroup having the smallest NRL. The effect of NRL shortening in cancer reported here may be not limited to CLL. In a separate work, we found a similar NRL shortening effect in paired tumour versus normal breast tissues from breast cancer patients (Jacob D.R., Guiblet W.M., Mamayusupova H., Shtumpf M., Gretton S., Correa C., Dellow E., Ruje L., Ciuta I., Agrawal S.P., Shafiei N., Vainshtein Y., Clarkson C.T., Thorn G.J., Sohn K., Pradeepa M.M., Chandrasekharan S., Klenova E., Zhurkin V.B. and Teif V.B., *under review*). Thus, NRL changes may be generally informative to identify tumour subtypes that are distinct in their disease course.

Our analysis of local changes of nucleosome occupancy showed that DNA methylation correlated with more stably bound nucleosomes, while nucleosomes that shift in CLL are at less methylated sites (Figure 2A-B). This relation was also observed in our studies in mouse embryonic stem cells (Teif et al. 2014; Wiehle et al. 2019). Such nucleosome repositioning is likely happening as an active process, which is reflected by the enrichment of histone deacetylases HDAC1 (Figure 2L) and chromatin remodellers BRG1 (SMARCA4) and CHD1 (Figures 2I-K). BRG1 is known to be involved in establishing B-cell identity (Bossen et al. 2015) and B-cell proliferation and germinal centre formation depends on enhancer activation by BRG1 (Schmiedel et al. 2021). Mutations in CHD1 homolog CHD2 are among the most frequent mutations in CLL and characteristic of the M-CLL subtype (Rodriguez et al. 2015). It is of note that this interplay between differential DNA methylation and nucleosome remodelling could also be one of the mechanisms that lead to NRL changes (Figure 1E).

Regions that undergo nucleosome occupancy changes were particularly enriched for CTCF binding sites, enhancers and promoters, and changes at promoters were associated with CLL-specific and immunity-related pathways (Figure 3). On a larger scale, we observed changes of nucleosome occupancy at enhancers and active chromatin compartments (Figure 4). Sites with CTCF binding loss become covered by nucleosomes and gain methylation in CLL subtype-dependent manner (Figure 5A-B). Among the seven histone modifications that we profiled, the most dramatic changes were correlated with the active enhancer mark H3K4me1 and the heterochromatin marks H3K9me3 and H3K27me3 (Figure 5C).

The analysis of the interplay between TF binding and nucleosome positioning allowed to reconstruct CLL-specific TF networks (Figure 6). The most enriched TF binding sites inside regions with differential nucleosome occupancy were related to immunity (such as IRF8 and IRF1), and specifically the germinal centre establishment (BCL6) (Capello et al. 2000; Mlynarczyk et al. 2019), as well as 3D genome organization (CTCF and its interaction partners, including cancer-specific competitor BORIS (CTCFL) and methylation-dependent factor NRF1) (Domcke et al. 2015). Importantly, several deregulated TFs detected in this study have not been previously associated with CLL, although some are involved in other cancers (Figure 6). Overall, three major groups of CLL-specific TFs were identified: (i) TFs for which nucleosome occupancy profiles around binding sites had quantitatively similar change in the average occupancy level in U-CLL and M-CLL versus NBCs, including PAX9 and PAX5 (Figure 6B). Interestingly, higher expression of PAX9 was reported for U-CLL and linked to poorer clinical outcome (Rani et al. 2017), whereas PAX5 is critical for both non-malignant B-cell development and CLL pathogenesis and evolution (Puente et al. 2015; Ott et al. 2018; Klintman et al. 2021). (ii) TFs for which nucleosome profiles in M-CLL were closer to NBC, such as TCLF5, HES5 and E2F3 (Figure 6B). E2F3 is a miR-34a target and a repressor of the ARF/p53 pathway. miR-34a has been linked to the chemotherapy resistance network in CLL and its low expression is associated with p53 aberrations and Richter transformation (Balatti et al. 2018). (iii) TFs with different shape of nucleosome occupancy profiles between CLL and NBCs, such as a depletion of nucleosome occupancy in NBCs changed to a peak in CLL (e.g., JUND, PKNOX2 and FOSL1 in Figure 6H-J). PKNOX2 acts as tumour suppressor in gastric cancer via activation of IGFBP5 and p53 (Zhang et al. 2019). JUND is involved in diffuse large B-cell lymphoma (Papoudou-Bai et al. 2016), while FOSL1 has been reported to promote carcinogenesis and metastasis in various cancer types (Jiang et al. 2020; Zhang et al. 2021), but not in CLL. FOSL1 gene encodes the FRA1 antigen (Jiang et al. 2020), which forms an AP-1 complex with the protein of the JUN family to exert its oncogenic effect. Thus, it is likely no coincidence that both FOSL1 and JUND change the shape of nucleosome occupancy profiles at their binding sites. Gene expression of FOSL1 is decreased ∼2-fold in CLL versus NBCs, which may explain why many of the binding sites occupied by FOSL1 in NBCs become occupied by a nucleosome in CLL. It is also worth noting that AP-1 is known to interact with SWI/SNF chromatin remodellers (Vierbuchen et al. 2017), thus providing possible connection to the involvement of BRG1 in active nucleosome repositioning (Figure 2I-J).

Finally, we asked whether nucleosome positioning *per se* can be used as a biomarker for CLL patient stratification. Principal component analysis (PCA) of nucleosome occupancies in regions that lost or gained nucleosomes in U-CLL versus NBCs allowed identification of not only U-CLL and NBCs, but also M-CLL, which was not included in the initial selection of nucleosome-sensitive regions (Figure 7C-F). A similar analysis based on gene expression (Figure 7A) was less effective. In part, this can be explained by the fact that CLL-specific gene expression is mostly occurring in lymph nodes – the primary site of CLL proliferation – while B-cells from peripheral blood are already “ primed” for malignant-specific expression by nucleosome positioning, but need in addition a specific microenvironment for this expression to occur (Herishanu et al. 2011). Furthermore, PCA based on DNA methylation was confounded by the developmental trajectory of individual CLL cases (Figure 7B), while nucleosome occupancy-based analysis was able to resolve this issue, e.g., when using subsets of regions which lost nucleosomes in CLL. Thus, given the extensive heterogeneity in CLL, the analysis of nucleosome positioning could enhance our understanding for critical malignant transformation events that are subtype-specific, indicative for distinct cell of origin and diverse microenvironmental cues.

In summary, nucleosome positioning undergoes large-scale changes in tumour cells, as reflected in the NRL decrease in CLL in addition to specific local changes at functional genomic elements. Thus, the analysis of aberrant nucleosome positions in primary tumour cells is a powerful approach for distinguishing malignant subtypes and, at the same time, informs about associated deregulation mechanisms. One particular area of application of the nucleosome positioning framework described here is patient diagnostics based on cfDNA, using liquid biopsies of peripheral blood samples. In the latter case, even if the condition that is being diagnosed is a solid tumour, the majority of cfDNA comes from nucleosome-protected regions in blood cells, similar to the nucleosome maps studied here. Finally, defining patient subsets using nucleosome positioning is a promising novel approach to stratify patient subgroups which can be exploited for personalized precision oncology.

## Methods

### Generation of normalised nucleosome occupancy profiles

The MNase-assisted histone H3 ChIP-seq reads in NBCs, M-CLL and U-CLL were from our previous study (EGA archive, access number EGAS00001002518) (Mallm et al. 2019). Paired-end sequenced reads were mapped with Bowtie to human genome hg19 allowing up to 1 mismatch and retaining only uniquely mappable reads. This resulted in ∼150 million paired reads/sample for a total of 26 samples (6 NBCs, 16 M-CLL and 4 U-CLL) (Supplemental Table S6). The cfDNAtools script extract_nuc_sizes.pl developed here (https://github.com/TeifLab/cfDNAtools) was used to extract fragments of size 120-180 bp, after which we obtained on average 105 million paired reads/sample. NucTools (Vainshtein et al. 2017) was used to calculate genome-wide nucleosome occupancies per chromosome with single base pair resolution, which were normalised separately for each sample by the corresponding sequencing depth. Averaged profiles for a given sample were calculated based on the normalised profiles of individual patients using NucTools with 100-bp running window. The resulting genome-wide nucleosome occupancy maps for the complete sample set derived here are available at GEO under the access code GSE158745. Since we limited our analysis to uniquely mappable regions, realigning this data to a newer human genome assembly GRCh38 will not affect the major conclusions of this study.

### Identification of nucleosomes with changed location in CLL

The annotation as stable/shifted nucleosome refers to DNA fragments of 120-180 bp size, which had genomic coordinates overlapping >95% between all replicates of all conditions (“ stable”) or less than 80% overlapping between NBCs and CLL (“ shifted”). Only nucleosomes that were stable across all NBC samples (overlapped >95% between all NBC replicates) were included in this analysis. The nucleosomes that overlap >95% (shifted <5%) were determined by performing pairwise intersection between the corresponding samples using BedTools with parameters –u -f 0.95. Coordinates of nucleosomes that were shifted >20% were obtained by intersection with parameters -f 0.80 -r -v. Aggregate methylation profiles at such nucleosomes were calculated using NucTools considering only CpGs with beta-values ≥0.8, which were selected using script methylationThresholds.pl available at github.com/TeifLab/cfDNAtools.

### Identification of genomic regions of differential nucleosome occupancy in CLL

Differences in nucleosome occupancy between NBCs, M-CLL and U-CLL were identified with NucTools. First, we applied the NucTools bed2occupancy_average.pl script to average nucleosome occupancy values genome-wide with a sliding window of 100 bp or 1000 bp (as detailed below) separately for each sample. Next, we defined regions which have stable nucleosome occupancy within each condition using the stable_nucs_replicates.pl script. This step included the normalisation of nucleosome occupancy profiles by the sequencing depth per chromosome per sample. Then normalised profiles in each 100-bp window were averaged across all samples in a given condition (6 NBCs, 16 M-CLL, 4 U-CLL). Genomic regions where the relative error after averaging was <0.5 were termed “ stable-nucleosome” regions and included in the following analysis. These stable-nucleosome regions were used to perform pairwise occupancy comparisons between M-CLL vs NBCs, U-CLL vs NBCs and U-CLL vs M-CLL with the script compare_two_conditions.pl. This script calculates the relative occupancy change (O_diff_) as O_diff_ = 2 x (<O_N1_> - <O_N2_>)/(<O_N1_> + <O_N2_>). The parameter <O_N1_> corresponds to the averaged nucleosome occupancy in data set 1 and <O_N2_> is the average occupancy in data set 2. Regions that lost or gained nucleosomes between conditions are defined as those where the relative occupancy change exceeds a given threshold. A 1000-bp sliding window and O_diff_ = 0.70 were used to calculate relative occupancy changes for the comparison of U-CLL vs NBC and M-CLL vs NBC reported in Figure 7 and Supplemental Figures S8-S9. For all other calculations a 100-bp sliding window size was used with O_diff_ = 0.99 for U-CLL vs NBC and M-CLL vs U-CLL and O_diff_ = 0.70 for M-CLL vs NBC due to the lower number of regions reported.

### Enrichment analysis

Fold enrichment of genomic features in lost/gain nucleosome-regions were calculated using BedTools “ fisher” function (Quinlan 2014). Two tail p-value was reported along with the enrichment ratio. Pathway enrichment analysis was performed in DAVID 6.8 (Jiao et al. 2012) and gprofiler2 (Raudvere et al. 2019) and visualized with Origin Pro (OriginLab).

### NRL analysis

NRL was calculated with NucTools following our standard protocol described previously (Vainshtein et al. 2017), but discarding the first peak of the nucleosome-nucleosome distance distributions to make the calculation more robust against the level of MNase digestion. The standard error of the NRL estimation was <1 bp for all samples.

### CTCF binding analysis

CTCF binding sites inside CTCF ChIP-seq peaks were predicted with GimmeMotifs (Bruse and Heeringen 2018) using the weight matrix MA0139.1 from JASPAR (Khan et al. 2018), accepting motifs with similarity score >80%. These sites were intersected using BedTools with the experimentally determined CTCF ChIP-seq peaks for each sample from (GSE113336) to obtain CTCF binding sites specific to NBCs, M-CLL and U-CLL. CTCF sites present in both NBCs and B-cells from all CLL patients where termed “ common”, and sites present only in NBCs but not in CLL were termed “ lost”. 41,202 CTCF-bound ChIP-seq peaks observed both in NBCs and CLL were defined as common, which contained 43,754 unique CTCF motifs. 718 CTCF-bound ChIP-seq peaks that were lost in CLL vs NBCs contained 708 CTCF motifs. 430 peaks containing 124 motifs which gained CTCF binding in M-CLL vs NBCs; 369 peaks containing 255 motifs gained CTCF binding in U-CLL vs NBCs). Additionally, the computationally predicted CTCF motifs were split into three quantiles based on the motif similarity determined with TFBStools (Tan and Lenhard 2016). Sites with similarity score of 80-81% were defined as quantile 1 CTCF motifs, 81-83% as quantile 2 CTCF motifs and motifs with score higher than 83% were classified as quantile 3.

### Calculation of aggregate profiles of nucleosome occupancy and histone modifications

Calculation of aggregate nucleosome profiles was performed with HOMER (Heinz et al. 2010), using only DNA fragments with sizes of 120-180 bp unless specified otherwise in the text. Aggregate profiles of seven histone modifications measured with ChIP-seq for this cohort (Mallm et al. 2019) were calculated using HOMER considering all ChIP-seq reads uniquely mapped with Bowtie to hg19 genome with up to 1 mismatch. Calculation of aggregate methylation profiles was done using custom Perl scripts as detailed below.

### Calculation of aggregate DNA methylation profiles

We calculated two types of DNA methylation profiles. In the first type of analysis (Figures 2A-B, 2E-H, 5A-B, 7B, S4, S5, S7), CpGs with beta values higher than 0.8 for each sample were extracted using a script methylationThresholds.pl available at github.com/TeifLab/cfDNAtools, split into chromosomes using NucTools script extract_chr_bed.pl followed by NucTools script bed2occupancy_average.pl to calculate DNA methylation density arising from such CpGs genome-wide with sliding window size 1-bp. This was then used for calculating occupancy profiles around genomic features with NucTools script aggregate_profile.pl. In this type of calculation, each CpG with beta-value >0.8 contributes equally. In the second type of calculation (Figures 4A-D), DNA methylation data reported previously for this cohort (Mallm et al. 2019) was processed with the Perl script bed2occupancy.v3d.methyl.pl available at github.com/TeifLab/cfDNAtools as in (Wiehle et al. 2019), summing up the actual methylation beta-values (which take values in the interval [0, 1]) for each individual CpG located at a given distance from the genomic feature of interest. When the corresponding graph refers to “ DNA methylation (a.u.)”, this corresponds to the value obtained by summation of all corresponding beta-values without further normalization. DNA methylation profiles of individual examples regions were calculated considering all methylation beta-values (Figures 1D, S3).

### Principal component analysis (PCA)

DNA methylation and nucleosome occupancy values averaged over a 100-bp window for each sample were intersected with coordinates of genomic features of interest (e.g., promoters or regions that lost/gained nucleosome occupancy in CLL) to create a matrix of occupancy values at the specified genomic regions, for each sample. Principal component analysis was performed in R and the relationship of principal components 1 and 2 was visualised in OriginPro (originlab.com).

### Annotation of genomic features

The coordinates of 100,000-bp genomic regions, annotated as A and B chromatin compartments in NBCs from peripheral blood, were kindly provided by Vilarrasa-Blasi et al. (Vilarrasa-Blasi et al. 2021) as BED files in hg38 genome assembly. This included 8,300 A- and 8,786 B-compartments in NBCs. LiftOver was used to convert the coordinates from hg38 to hg19, with default options. Conversion failed on 48 and 35 A and B compartment records, respectively. Promoters were defined as regions +/- 1,000 bp around TSS, based on RefSeq annotation. Enhancers specific to NBCs and CLL were defined based on the same cohort following our previous report (Mallm et al. 2019). Binding sites of SP1 and Chd1 were determined based on ChIP-seq peaks in lymphoblastoid cell line GM12878 reported by the ENCODE consortium (GEO entries GSM803363 and GSM935301 correspondingly). Binding sites of HDAC1 in peripheral blood mononuclear cells from CLL patients with 11q deletion genotype were determined based on ChIP-seq peaks downloaded from GEO entry GSE216287. Enrichment of BRG1 was calculated using ChIP-seq of BRG1 binding in naïve B-cells J1 (GSM971343) (Abraham et al. 2013). For the latter, we mapped the raw data to the hg19 genome, calculated aggregate profiles with HOMER, then normalized these by dividing the BRG1 ChIP-seq signal by the corresponding Input reported by the authors.

## Data access

The processed sequencing data generated in this study have been submitted to the NCBI Gene Expression Omnibus (GEO; https://www.ncbi.nlm.nih.gov/geo/) and will be made openly available upon the journal publication. The raw sequencing data including MNase-assisted H3 ChIP-seq are available at EGA (EGAS00001002518).

## Acknowledgements

This work was supported by the Wellcome Trust grant 200733/Z/16/Z and Cancer Research UK grant EDDPMA-Nov21\100044 to VBT, the Genetics Society grant to KVP and VBT, DFG grants for subprojects B1, B2 and Z1 within SFB1074 to DM, SS and KR, and the Leukaemia & Lymphoma NI R2740CEM and the Academy of Medical Sciences SBF005\1113 grants to EK. VT thanks Victor Zhurkin for fruitful discussions.

## Author contributions

KVP performed analyses related to CTCF, regulatory regions, remodellers, A/B compartments, histone modifications and DNA methylation, as well as PCA analysis for patient stratification. CMD, CX, CTC and CF performed analyses related to TF binding. JPM contributed to the data preprocessing. LR performed data visualisation. LCK performed patient classification based on B-cell developmental status. YV maintained and upgraded the NucTools software. VBT performed initial data analyses leading to this manuscript and NRL calculations. SS and DM advised on medical aspects. EK, KR and VBT supervised the study. VBT and KR wrote the manuscript draft. All authors read and approved the final manuscript.

## Code availability

Our results make use of published software tools with detailed parameters included in the Methods. Additional custom scripts used to generate these results are available on GitHub at https://github.com/TeifLab/cfDNAtools.

