## Supplementary Materials for "Nucleosome repositioning in chronic lymphocytic leukaemia"

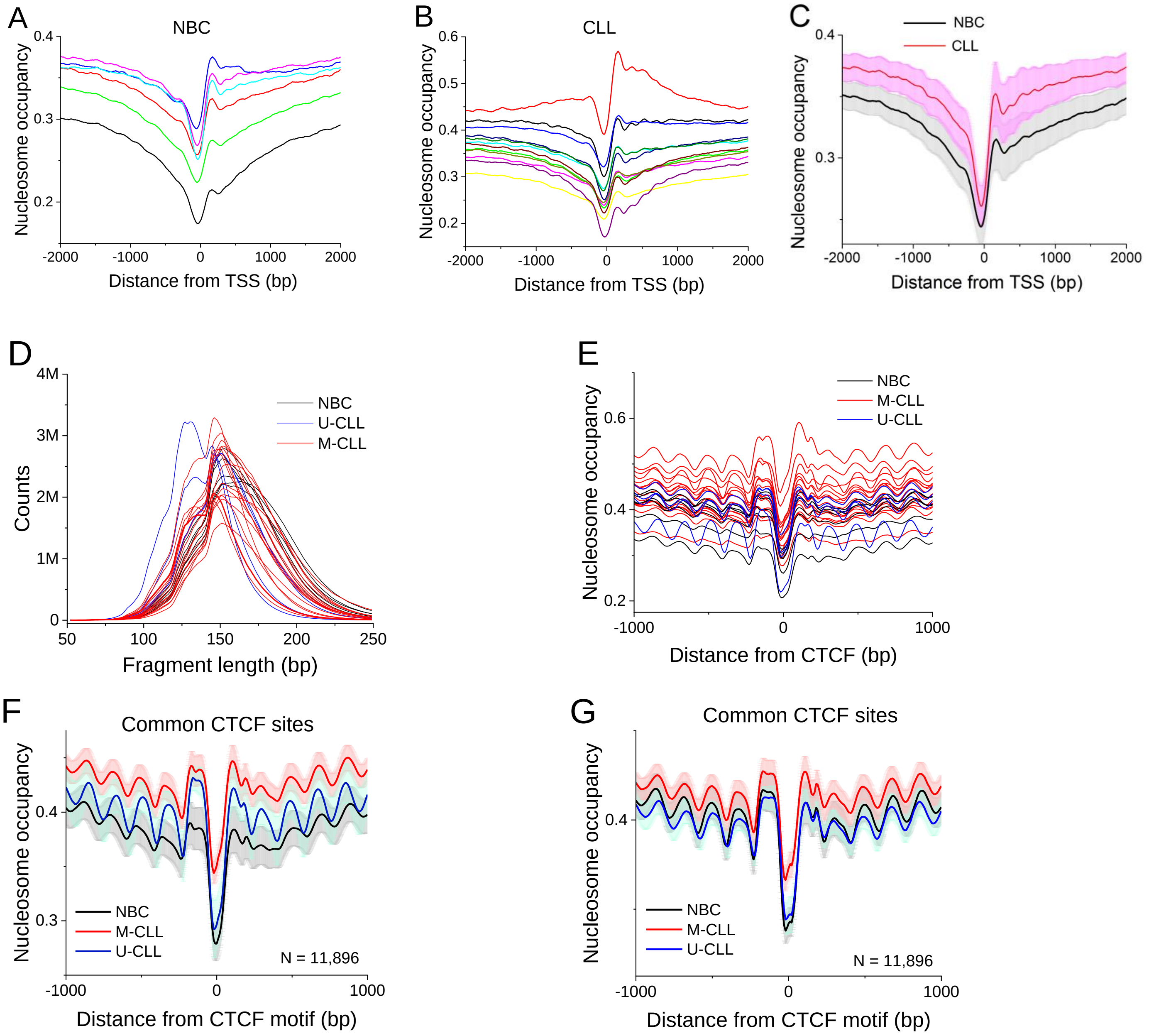


**Figure S1. Aggregate profiles and fragment size distributions for individual samples as well as averaged within each condition.** (A-C) Average nucleosome occupancy profiles around the TSS. (**A**) NBC nucleosome occupancies calculated for each replicate experiment. (**B**) Same as panel B for CLL B-cells. (**C**) NBCs (black) and CLL B-cells (red). All DNA fragment sizes were considered. (D-G) Distribution of nucleosomal fragment lengths and nucleosome occupancy around bound CTCF. (**D**) Distribution of nucleosomal DNA fragment lengths across all samples. NBC, black; M-CLL, red; U-CLL, blue. (**E**) Nucleosome occupancy profiles around common DNA sequence motifs that are bound by CTCF in all B-cell samples from CLL patients and NBC individuals. (**F**) Averaged nucleosome occupancy profiles taking into account all nucleosomal DNA fragment sizes. The standard error of averaging between samples is shown for each line in light colours. The number of regions (N) is indicated on the graphs. (**G**) Same as panel F but taking into account only DNA fragments with sizes 120-180 bp. The grey/pink/blue areas show the corresponding standard errors of averaging.

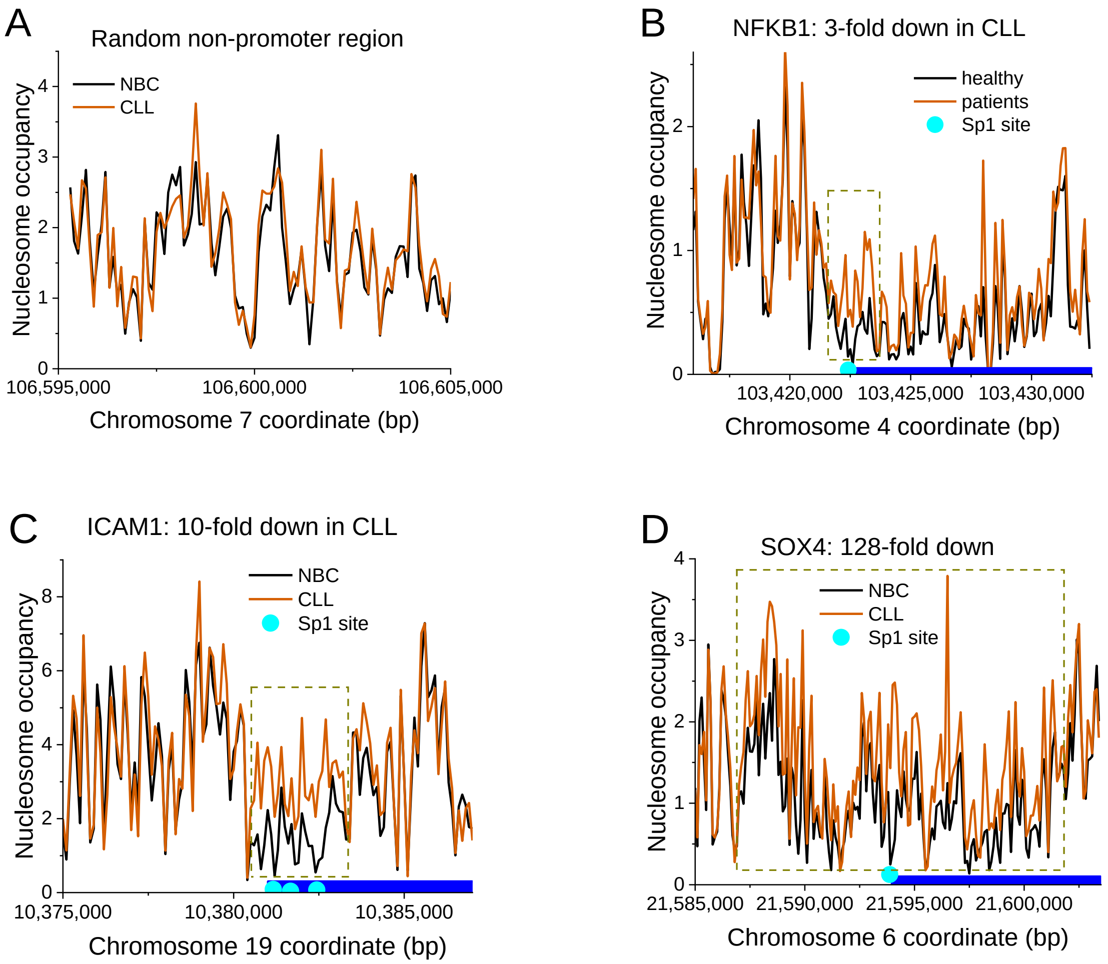


**Figure S2. Nucleosome occupancy profiles in exemplary genomic regions**. Data were averaged over all NBCs (black) and all CLL (brown) samples. Blue lines indicate gene locations. Light blue circles indicate Sp1 binding sites in the lymphoblastoid cell line GM12878. All DNA fragment sizes were considered. (**A**) Randomly selected non-regulatory region. (**B**) NFkB1 promoter. (**C**) ICAM1 promoter. (**D**) SOX4 promoter.

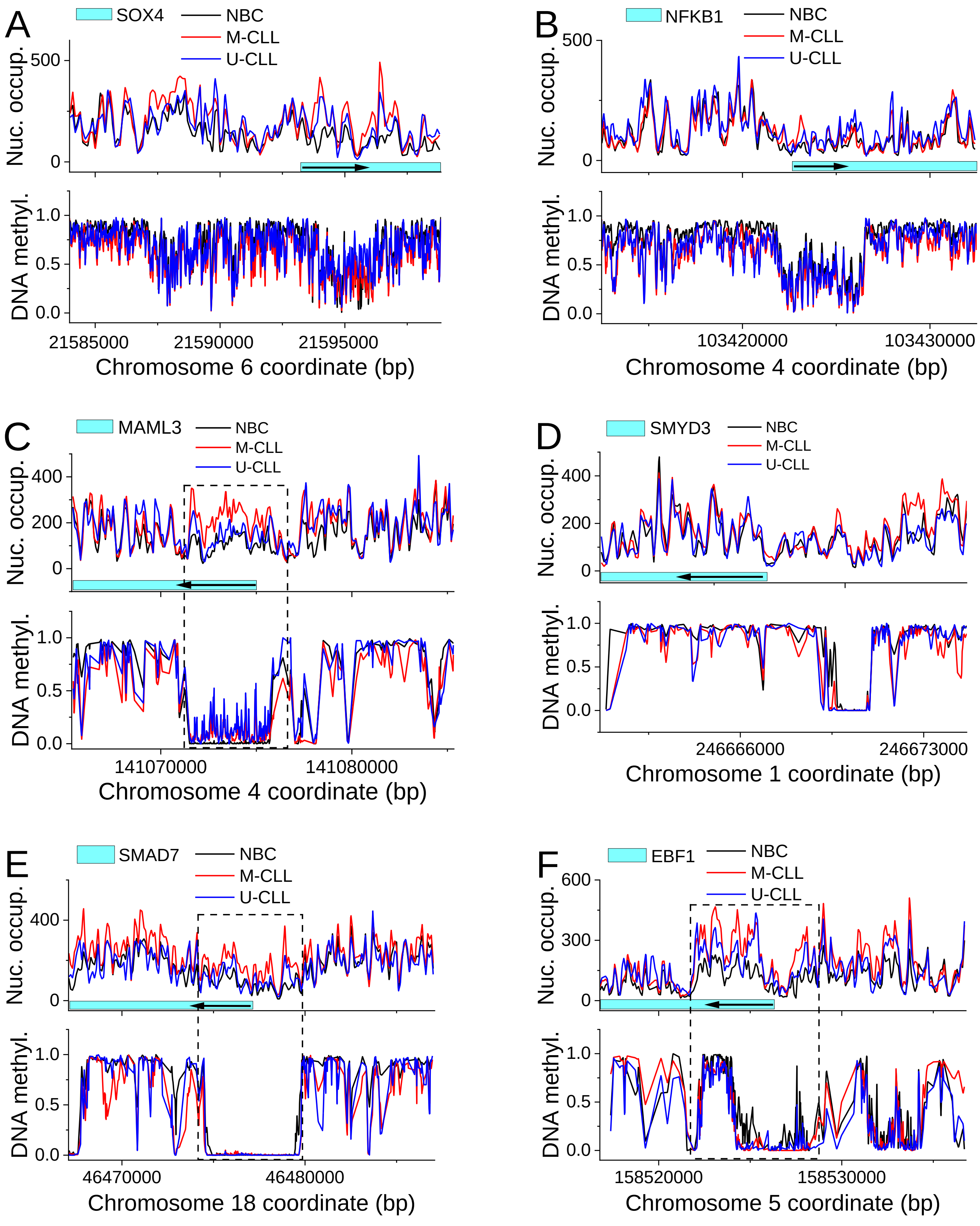


**Figure S3. Nucleosome occupancy and DNA methylation (bottom) profiles around exemplary promoters.** Averaged profiles were computed for DNA fragment sizes between 120 and 180 bp. NBC, black; M-CLL, red; U-CLL, blue. (**A**) SOX4. (**B**) NFKB1. (**C**) MAML3. (**D**) SMYD3. (**E**) SMAD7. (**F**) EBF1.

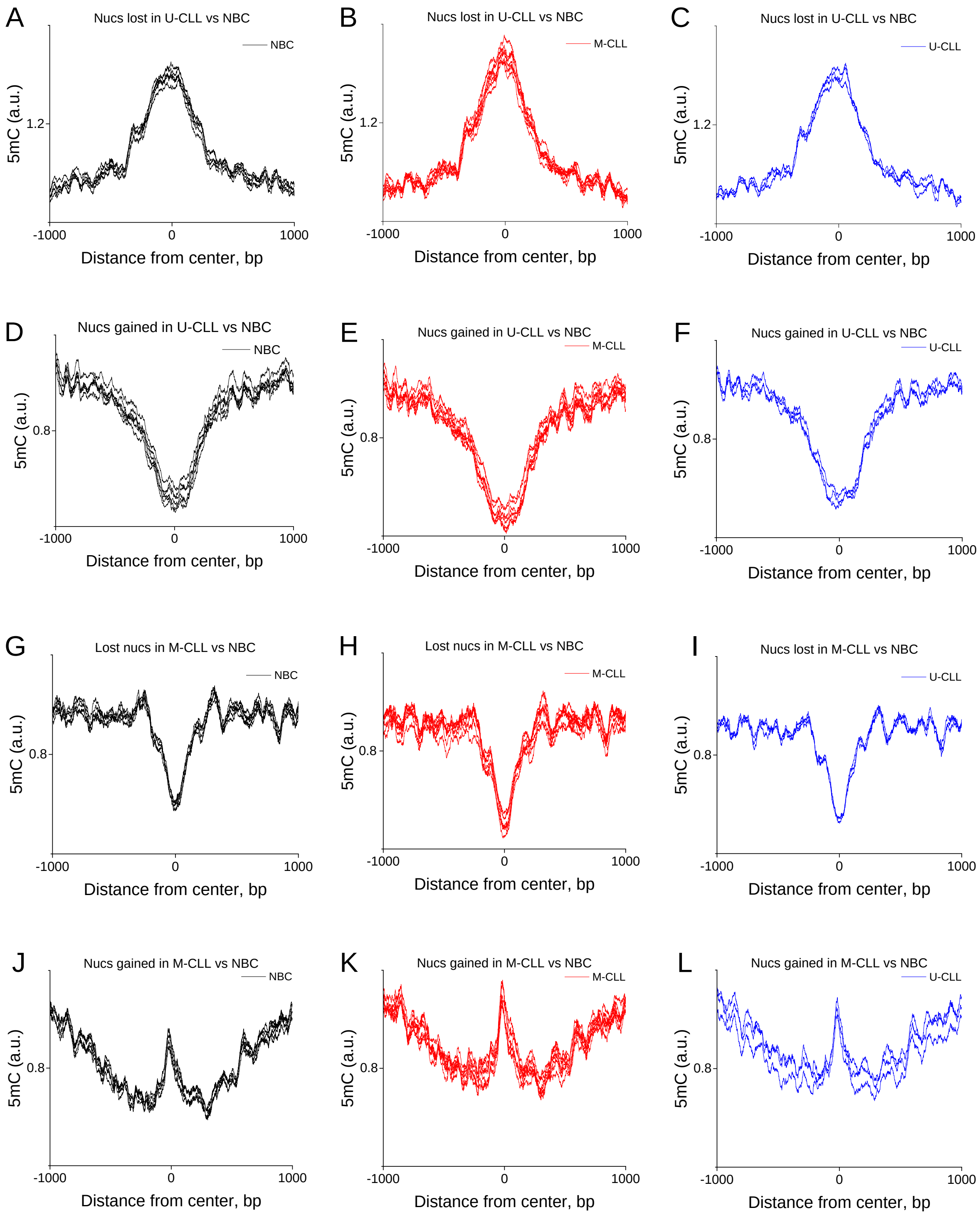


**Figure S4. Averaged DNA methylation profiles for the individual samples around centres of 100-bp regions that lost or gained nucleosomes in M-CLL or U-CLL vs NBCs.** The same coordinates of lost-nucleosomes and gained-nucleosomes regions are used as in Figure 2E-F, but here the profiles of DNA methylation were calculated for each individual sample without averaging across samples. Each sample is shown as a separate curve. The samples are grouped on the graph based on the medical including NBCs (black, left column), M-CLL (red, middle column) and U-CLL (blue, right column)**. First row:** DNA methylation around centres of 100-bp regions that lost nucleosomes in M-CLL vs NBCs (analogous to Figure 2E). **Second row:** the same for gained nucleosomes in M-CLL vs NBCs (analogous to Figure 2F). **Third row:** the same for lost nucleosomes in U-CLL vs NBCs (analogous to Figure 2G). **Bottom row**: the same for gained nucleosomes in U-CLL vs NBCs (analogous to Figure 2H).

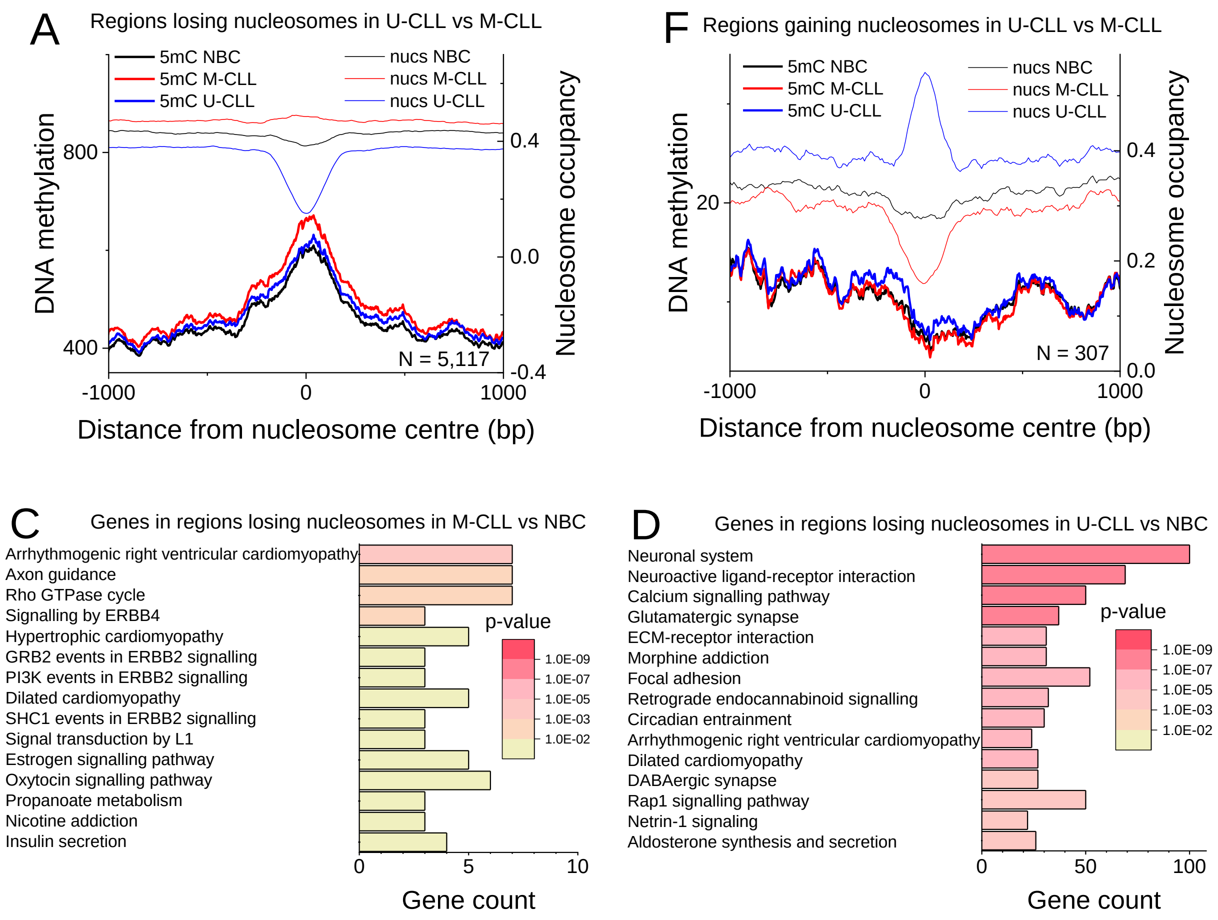


**Figure S5. Characterisation of genomic regions with lost/gained nucleosomes.** (A-B) DNA methylation profiles around regions with lost/gained nucleosomes. Averaged nucleosome occupancy (thin lines) and DNA methylation profiles (thick lines) were computed around the centres of 100-bp regions. The profiles are averaged over all NBC (black), M-CLL (red) and U-CLL (blue) samples. The number of regions (N) is indicated in the figure. (**A**) Regions with lower nucleosome occupancy in U-CLL vs M-CLL. (**B**) Regions with higher nucleosome occupancy in U-CLL vs M-CLL. (C-D) Gene ontology analysis of genes overlapping with regions which lost nucleosomes in CLL. Only DNA fragments with sizes 120-180 bp were considered. (**C**) Genes which overlap with regions that lost nucleosomes in M-CLL vs. NBC. (**D**) Genes which overlap with regions that lost nucleosomes in M-CLL vs. U-CLL.

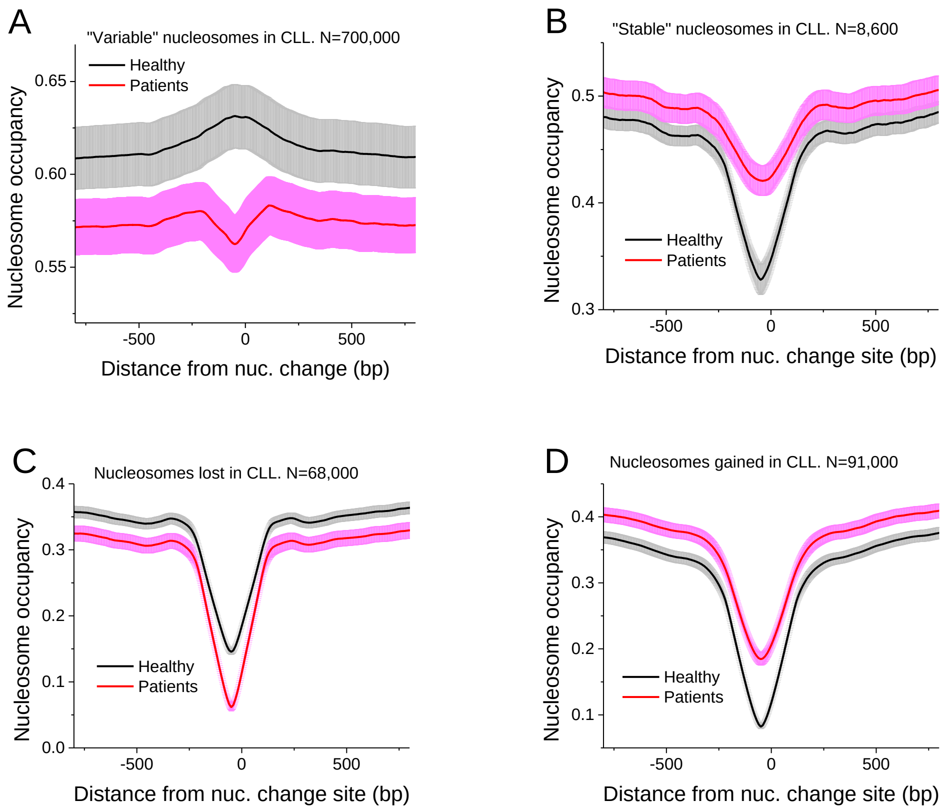


**Figure S6. Different classes of nucleosomes repositioned in CLL versus NBCs.** Nucleosome occupancy profiles were averaged over all NBC (black) and CLL (red) samples for all DNA fragments without size filtering. (**A**) Regions where nucleosome occupancy is variable across CLL patients while it is stable across NBCs. (**B**) Regions where nucleosome occupancy is stable in CLL while it is variable in NBCs. (**C**) Regions where average nucleosome occupancy in CLL decreases in comparison with NBCs ("lost nucleosomes"). (**D**) Regions where average nucleosome occupancy in CLL increases in comparison with NBCs ("gained nucleosomes")..

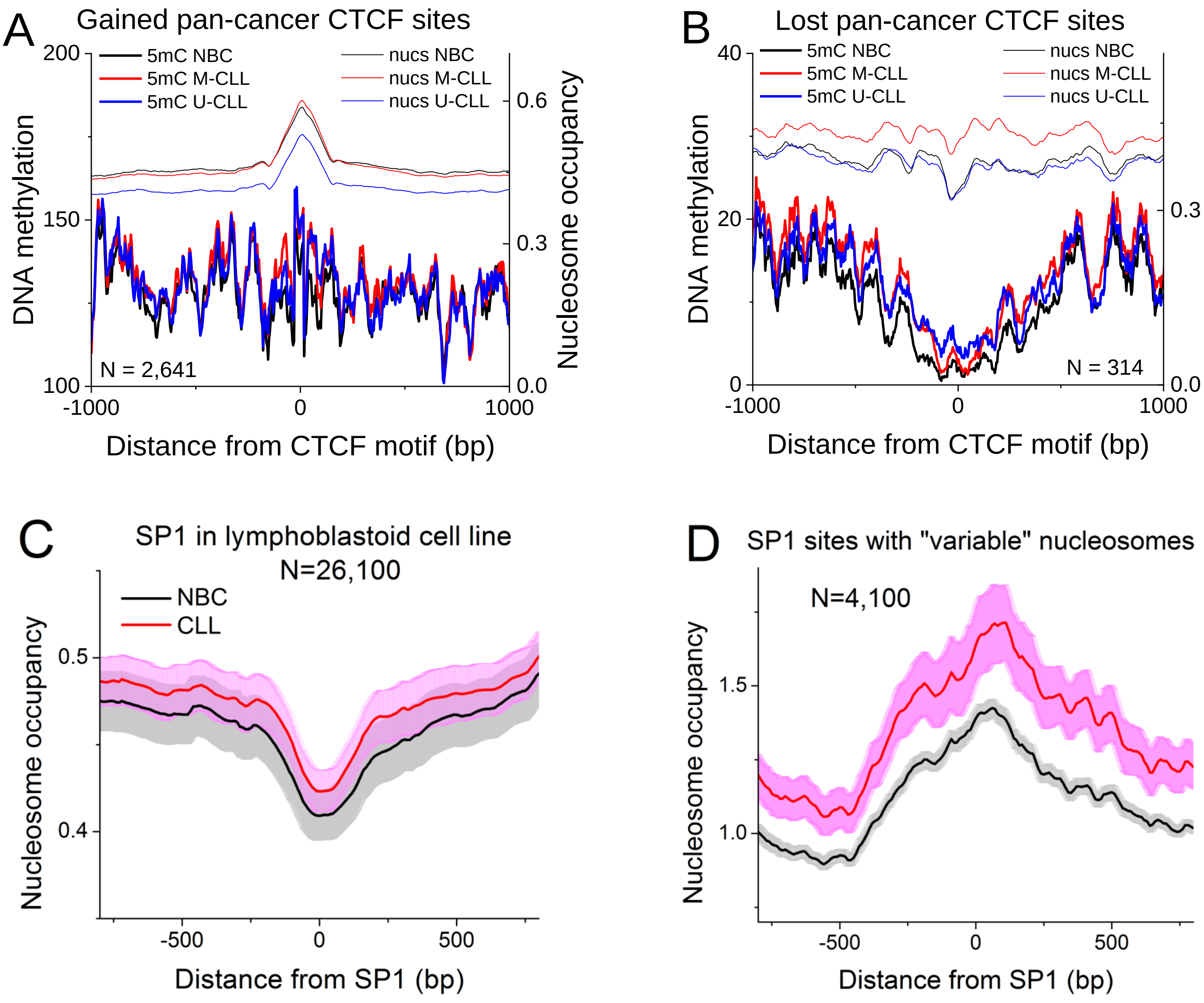


**Figure S7. Nucleosome occupancy determined in this study plotted around TF binding sites experimentally determined in previous publications**. (A-B) Nucleosome occupancy and DNA methylation around binding sites of the "pan-cancer dataset" of CTCF sites lost or gained across six cancers [Fang, *et al.* (2020) *Genome Biology* 21, 247]. The profiles were averaged over all NBC (black), M-CLL (red) and U-CLL (blue) samples. The number of regions (N) is indicated on the graph. (**A**) Averaged profiles around gained consensus CTCF sites. (**B**) Averaged profiles around lost consensus CTCF sites. (C-D) Averaged nucleosome occupancy profiles around binding sites of SP1 determined by ChIP-seq in lymphoblastoid cell line GM12878 from ENCODE (GSM803363). Nucleosome occupancy profiles averaged over all NBC (black) and CLL (red) samples around SP1 binding sites were calculated. (**C**) All SP1 binding sites in lymphoblastoid cell line GM12878. (**D**) Fraction of SP1 sites depicted in panel C that overlaps with regions that have variable nucleosome occupancy in CLL and stable nucleosome occupancy in NBCs.

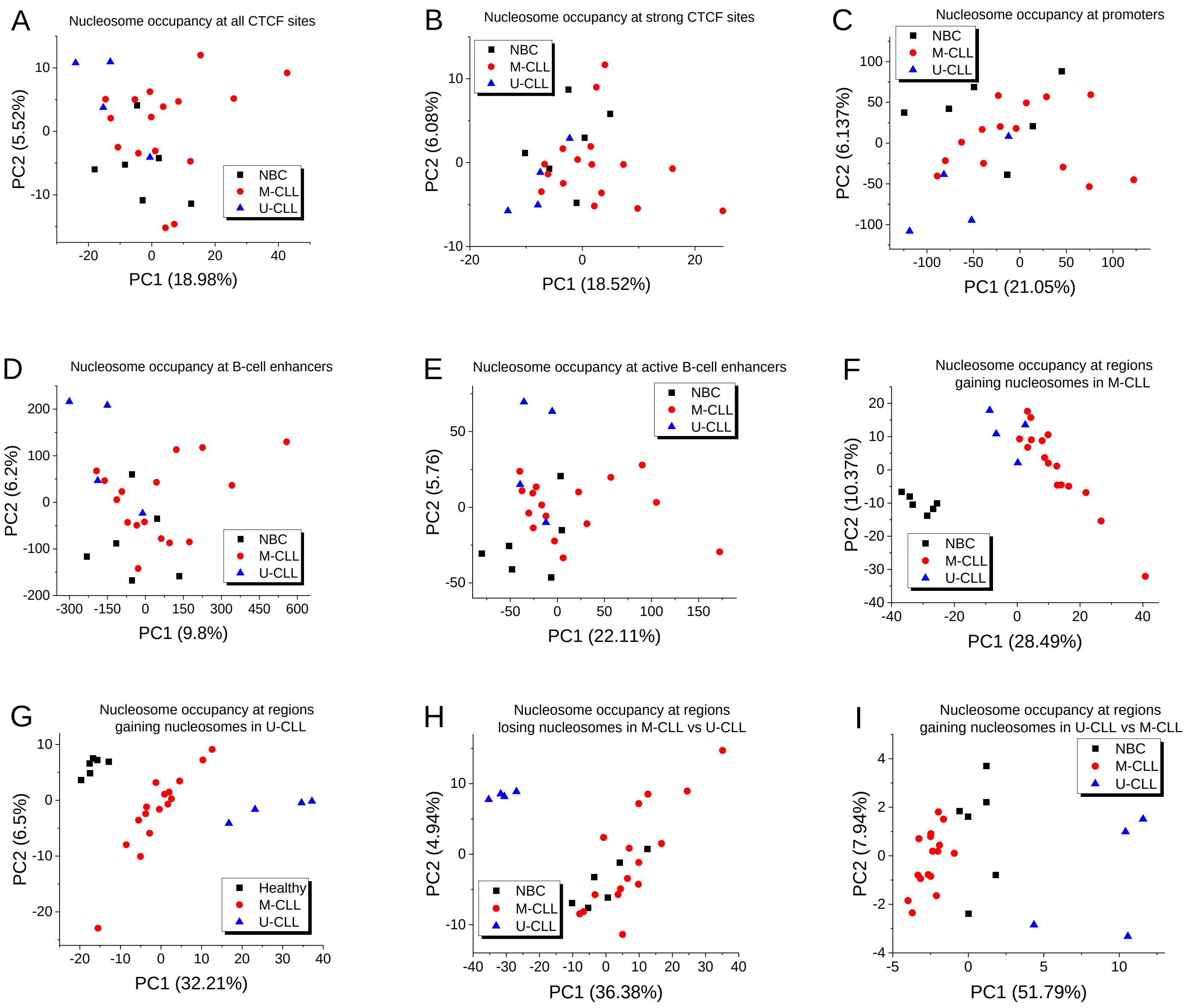


**Figure S8. Principal component analysis of nucleosome occupancy at genomic regions.** Each dot represents a replicate from NBCs (black), M-CLL (red) and U-CLL (blue). (**A**) CTCF all quantiles. (**B**) Quantile 3 CTCF sites. (**C**) Promoters. (**D**) B-cell enhancers. (**E**) Active B-cell enhancers. (**F**) regions with increased nucleosome occupancy in M-CLL. (**G**) Regions with increased nucleosomes occupancy in U-CLL, (H) regions with decreased nucleosome occupancy in U-CLL compared to M-CLL. (**I**) Regions with increased nucleosome occupancy in U-CLL compared to M-CLL.

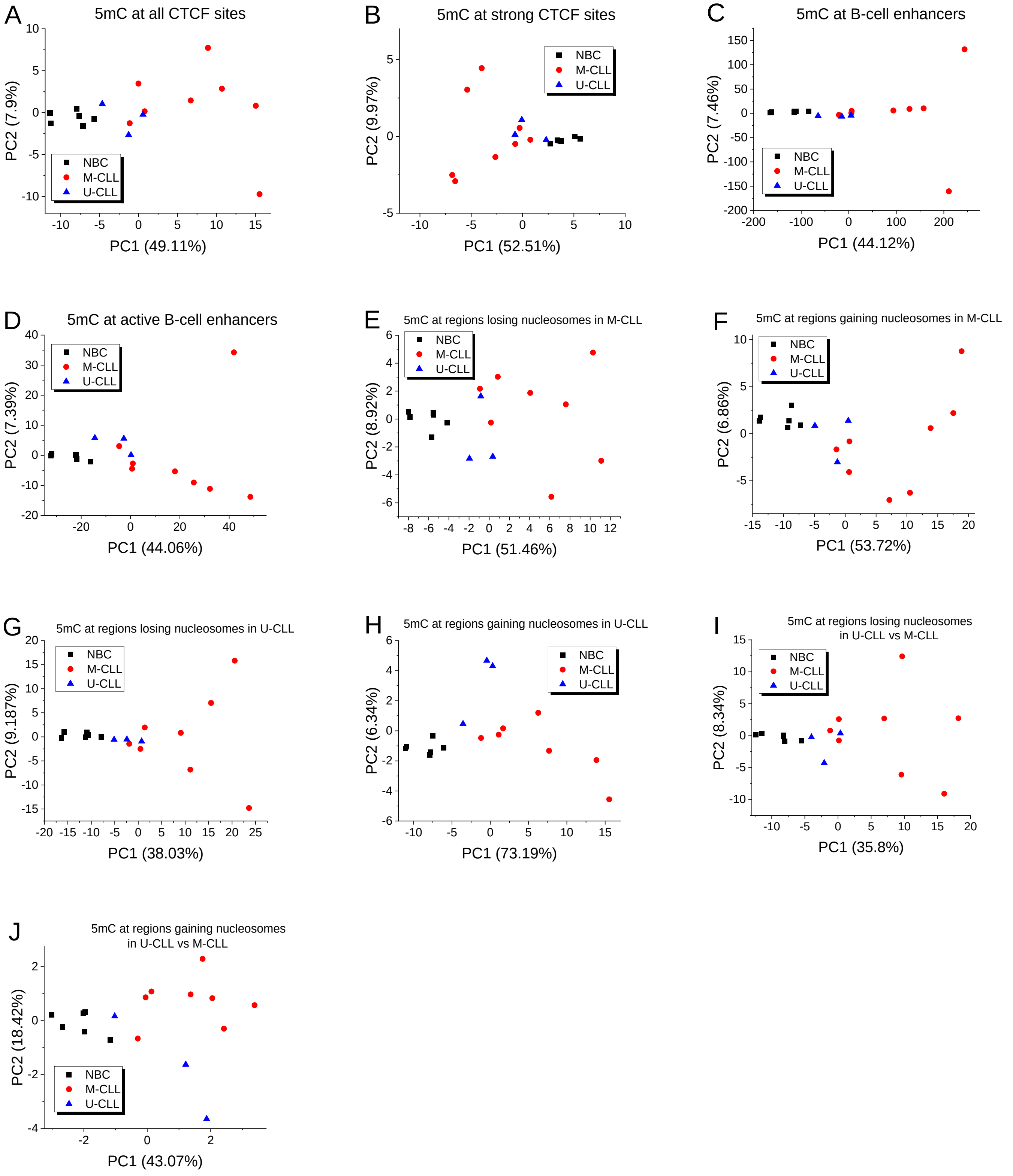


**Figure S9. Principal component analysis of DNA methylation occupancy at genomic regions.** Each dot represents a replicate from NBCs (black), M-CLL (red) and U-CLL (blue). (**A**) CTCF all quantiles. (**B**) Quantile 3 CTCF sites. (**C**) B-cell enhancers. (**D**) Active B-cell enhancers. (**E**) Regions with decreased nucleosome occupancy in M-CLL. (**F**) Regions with increased nucleosomes occupancy in M-CLL. (**G**) Regions with decreased nucleosome occupancy in U-CLL. (**H**) Regions with increased nucleosome occupancy in U-CLL. (**I**) Regions with decreased nucleosome occupancy in U-CLL compared to M-CLL. (**J**) Regions with increased nucleosome occupancy in U-CLL compared to M-CLL.

**Table S1**. Top pathways enriched among the genes whose promoters are marked by nucleosome gain in CLL versus NBCs based on all DNA fragments without fragment size-filtering.

| **Pathway name** | **# genes** | **P-value** | **Benjamini** |
|---|---|---|---|
| B cell receptor signaling pathway | 23 | 1.6e-5 | 3.0e-3 |
| T cell receptor signaling pathway | 26 | 3.2e-4 | 2.0e-2 |
| EGF receptor signaling pathway | 27 | 5.1e-3 | 1.1e-1 |
| Pancreatic cancer | 17 | 5.7e-3 | 1.2e-1 |
| Influenza Infection | 33 | 1.3e-3 | 2.0e-2 |
| -UTR-mediated translational regulation | 23 | 7.2e-3 | 8.6e-2 |
| CD40L Signaling Pathway | 7 | 7.5e-3 | 6.0e-1 |
| FGF signaling pathway | 25 | 1.0e-2 | 1.6e-1 |
| mTOR signaling pathway | 13 | 1.2e-2 | 2.2e-1 |

**Table S2**. Genes belonging to the B cell receptor signalling pathway (BCR) characterised by significantly increased nucleosome occupancy at their promoters in CLL versus NBCs.

<table>
<tr><th>Gene name</th><th>Expression (log2)</th><th>TF complex</th></tr>
<tr><td>BCL10</td><td>-0.981</td><td>-</td></tr>
<tr><td>CD79A</td><td>-0.897</td><td>-</td></tr>
<tr><td>CD79B</td><td>1.556</td><td>-</td></tr>
<tr><td>DAPP1</td><td>-0.250</td><td>-</td></tr>
<tr><td>JUN</td><td>-1.347</td><td rowspan="2">AP1</td></tr>
<tr><td>FOS</td><td>-4.770</td></tr>
<tr><td>MALT1</td><td>-0.0813</td><td>-</td></tr>
<tr><td>NFAT5</td><td>-0.474</td><td>-</td></tr>
<tr><td>NFATC1</td><td>-0.737</td><td>-</td></tr>
<tr><td>NFATC4</td><td>-0.437</td><td>-</td></tr>
<tr><td>NFKB1</td><td>-1.935</td><td rowspan="4">NFκB</td></tr>
<tr><td>NFKBIA</td><td>-2.865</td></tr>
<tr><td>NFKBIB</td><td>-1.715</td></tr>
<tr><td>NFKBIE</td><td>-1.972</td></tr>
<tr><td>PIK3CA</td><td>-1.437</td><td>-</td></tr>
<tr><td>PIK3CG</td><td>-1.330</td><td>-</td></tr>
<tr><td>PIK3R1</td><td>-0.655</td><td>-</td></tr>
<tr><td>PIK3R5</td><td>-2.504</td><td>-</td></tr>
<tr><td>RAC3</td><td>2.458</td><td>-</td></tr>
<tr><td>SYK</td><td>1.506</td><td>-</td></tr>
<tr><td>AKT2</td><td>-1.429</td><td>-</td></tr>
<tr><td>RAF1</td><td>-0.886</td><td>-</td></tr>
<tr><td>VAV1</td><td>0.649</td><td>-</td></tr>
</table>

**Table S3**. DNA sequence motifs in promoters of BCR genes marked by the nucleosome gain in CLL versus NBCs based on all DNA fragments without fragment size-filtering.

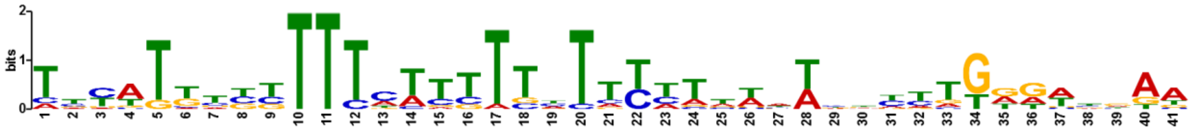


| Name | E-value | Width | # regions/total | Top 10 TFs recognizing this motif |
|---|---|---|---|---|
| Motif1 | 3.4e-032 | 41 | 15 / 23 | MTF1, FOXP1, IRF1, GATA6, ZFP105, FOXJ3, FOXP2, FOXK1, SOX11, SOX4 |

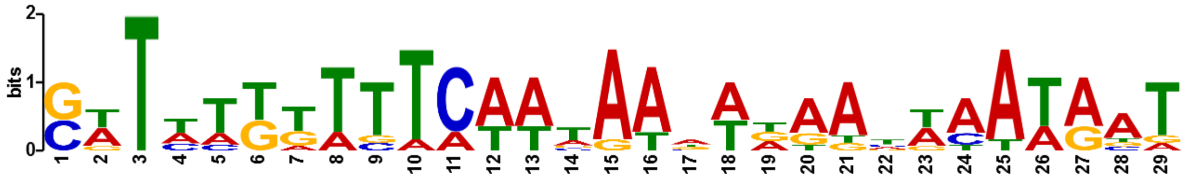


| Name | E-value | Width | # genes / total | Top 10 TFs recognizing this motif |
|---|---|---|---|---|
| Motif2 | 1.3e-011 | 29 | 9 / 23 | ELF3, DBX1, TBP, TCFAP2E, DBX2, HOXA4, SOX21, SOX1, LHX1, POU2F1 |

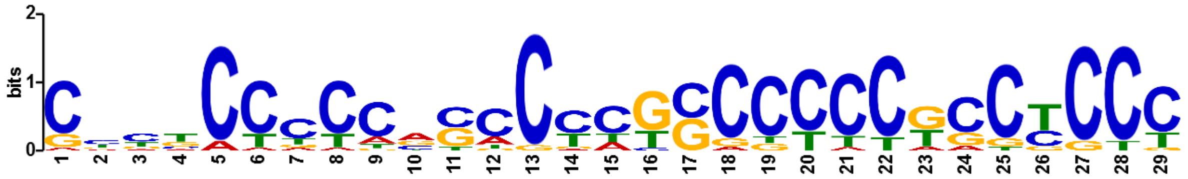


| Name | E-value | Width | # regions/total | Top 10 TFs recognizing this motif |
|---|---|---|---|---|
| Motif3 | 6.0e-011 | 29 | 21/23 | EGR1, SP1, SP2, ZNF263, EGR2, ZFP281, E2F3, KLF5, ZFP740, RREB1 |

The most common TF-binding motif inside nucleosomes gained at promoters in CLL versus NBCs based on all DNA fragments without fragment size-filtering:

<table>
<tr><td rowspan="2">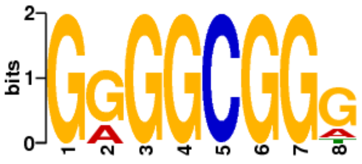
</td><td>Positives:<br>591/2508</td><td>P-value:<br>1.1e-37</td><td rowspan="2">Top TFs (P-values):<br>SP1 (4.2e-8), SP2 (4.4e-7), Zfp410 (1.2e-6), Bcl6b (1.3e-6), EGR1 (2.3e-6), E2F3 (5.2e-6), SP4 (7.4e-6), KLF7 (1.5e-5),<br>KLF5 (1.7e-5), E2F1 (4.0e-5)</td></tr>
<tr><td>Negatives:<br>255/2508</td><td>E-value:<br>4.5e-33</td></tr>
</table>

**Table S4. Nucleosome occupancy profiles around TF motifs inside ATAC-seq peaks lost in CLL.**

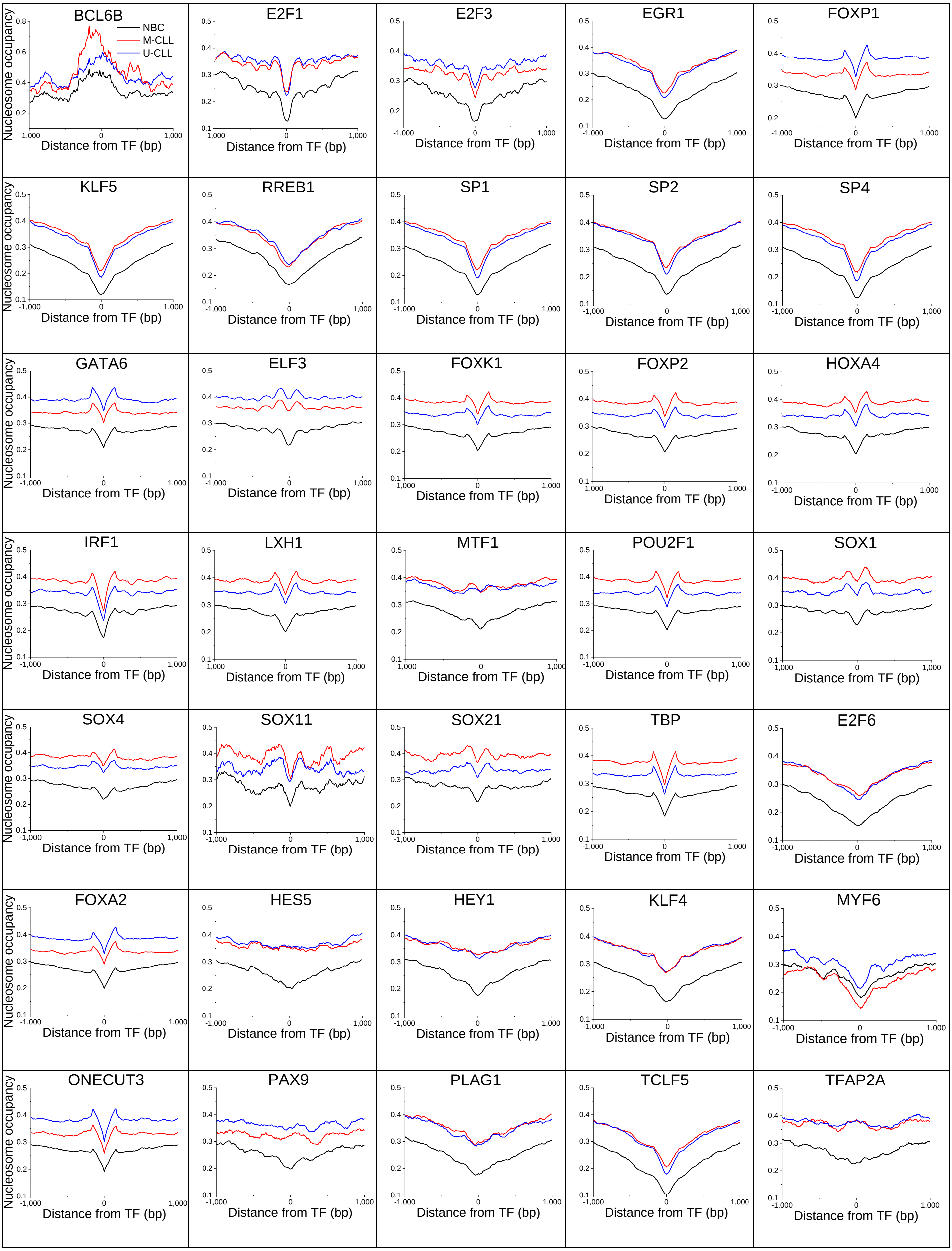

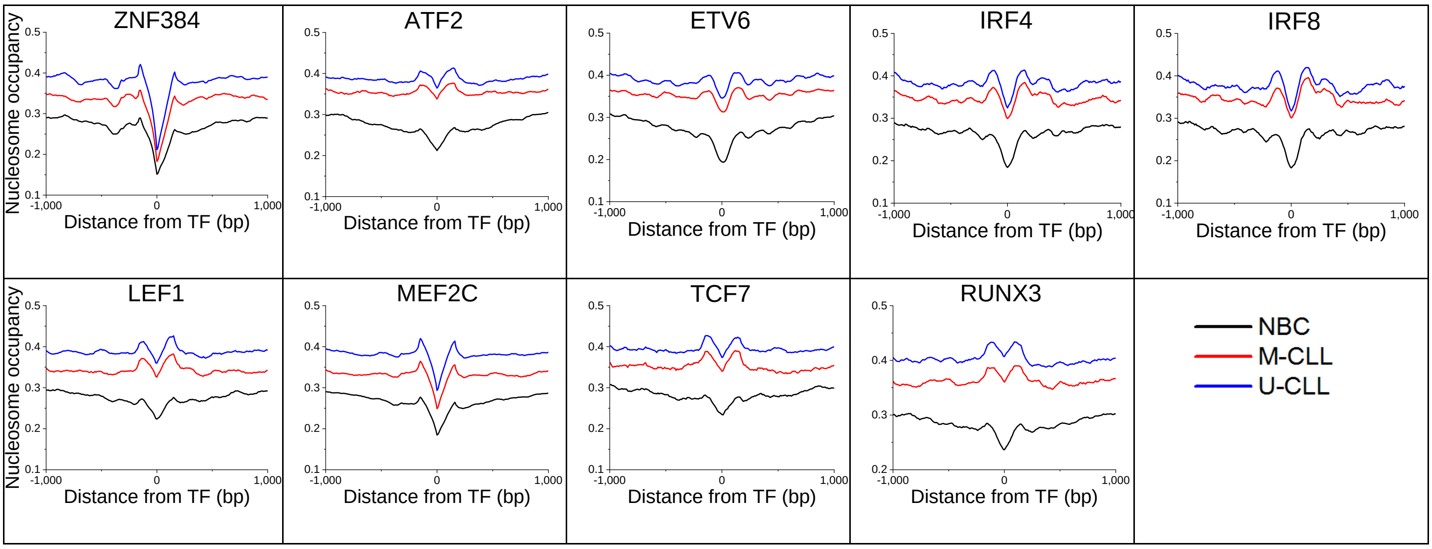
ZNF384
ATF2
ETV6
IRF4
IRF8
LEF1
MEF2C
TCF7
RUNX3
Nucleosome occupancy
Distance from TF (bp)
NBC
M-CLL
U-CLL

### Table S5. Nucleosome occupancy profiles around TF motifs inside ATAC-seq peaks gained in CLL.

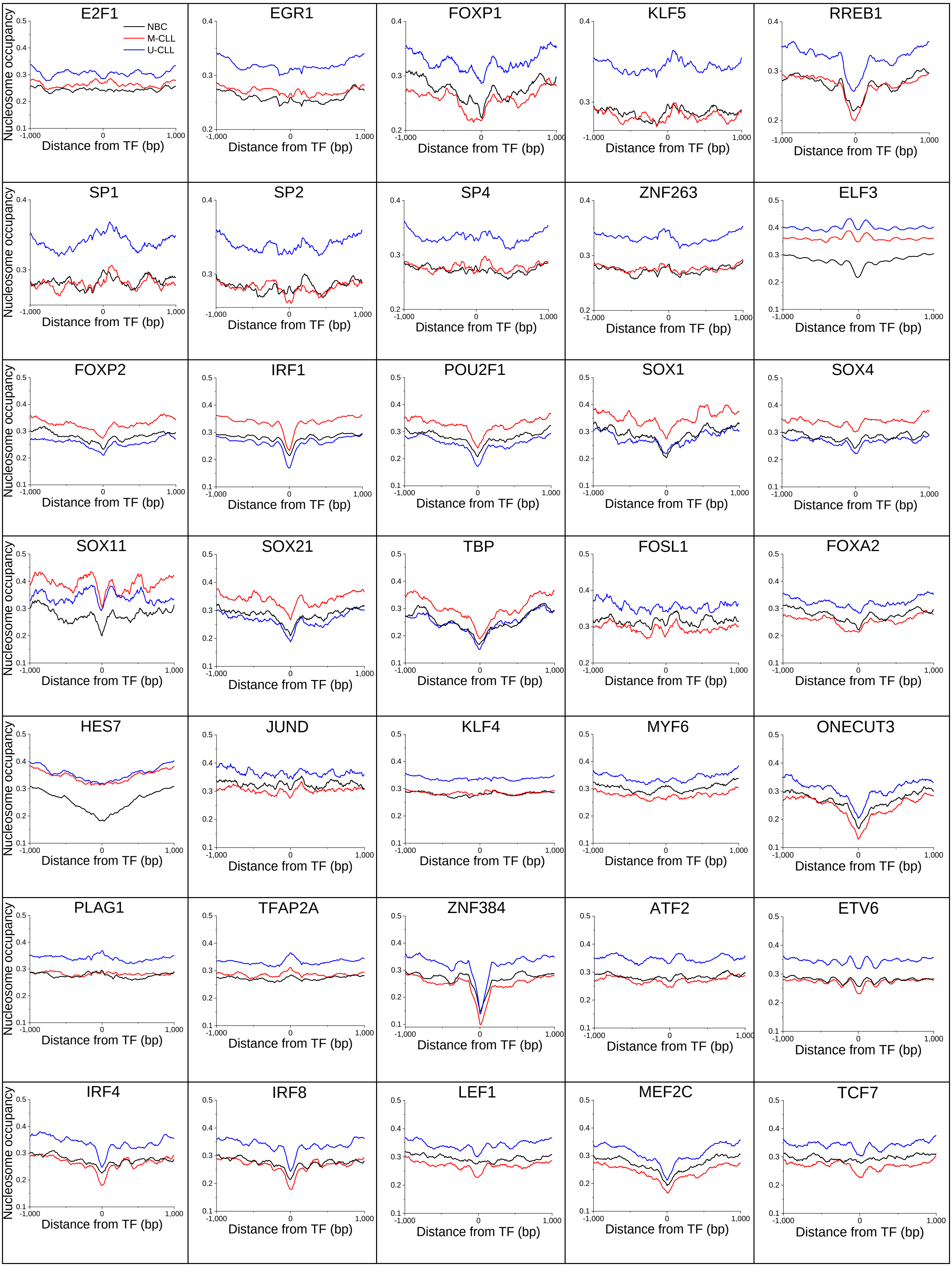

**Table S6. Numbers of mapped pared-end reads per sample obtained with MNase-assisted H3 ChIP-seq**

| Sample ID | Medical condition | Total reads | Uniquely mapped reads | Reads with 120-180 bp length |
|---|---|---|---|---|
| BK0916 | NBC | 243,395,806 | 156,114,070 | 115,084,219 |
| BK0925 | NBC | 213,127,910 | 152,130,702 | 98,398,076 |
| EM1161 | NBC | 243,736,807 | 165,241,426 | 99,882,287 |
| EM1170 | NBC | 234,925,334 | 162,500,056 | 104,669,238 |
| GN1125 | NBC | 252,089,321 | 162,210,501 | 115,898,081 |
| GN1134 | NBC | 250,370,079 | 163,532,966 | 113,564,829 |
| CX0577 | M-CLL | 180,397,845 | 124,541,409 | 89,461,451 |
| CX0586 | M-CLL | 148,542,561 | 90,417,857 | 66,256,250 |
| IO1203 | M-CLL | 250,008,531 | 172,563,637 | 118,455,189 |
| IO1212 | M-CLL | 227,423,203 | 115,555,639 | 82,220,448 |
| KL0603 | M-CLL | 199,514,059 | 152,428,741 | 113,813,980 |
| KL0612 | M-CLL | 201,491,142 | 156,937,195 | 117,823,099 |
| RC1143 | M-CLL | 247,411,232 | 154,569,836 | 103,538,797 |
| RC1152 | M-CLL | 249,953,057 | 150,406,896 | 91,005,433 |
| TH0447 | M-CLL | 152,342,103 | 102,373,391 | 83,419,384 |
| TH0456 | M-CLL | 208,423,556 | 164,154,393 | 133,526,874 |
| WW0529 | M-CLL | 238,814,095 | 160,835,696 | 118,360,090 |
| WW0538 | M-CLL | 248,025,727 | 169,421,890 | 127,499,614 |
| XP0962 | M-CLL | 145,359,217 | 105,872,205 | 84,439,719 |
| XP0970 | M-CLL | 137,340,079 | 102,328,977 | 82,145,439 |
| ZV1179 | M-CLL | 244,265,740 | 168,120,260 | 121,977,059 |
| ZV1188 | M-CLL | 203,569,432 | 144,209,193 | 96,795,047 |
| PP1093 | U-CLL | 211,561,008 | 151,806,790 | 113,488,851 |
| PP1102 | U-CLL | 217,337,789 | 155,733,690 | 121,070,657 |
| QU1019 | U-CLL | 179,948,637 | 132,369,587 | 109,207,547 |
| QU1028 | U-CLL | 208,754,718 | 149,923,687 | 114,946,538 |
| **Total:** | | **5,538,128,988** | **3,786,300,690** | **2,736,948,196** |
